# A single-cell atlas reveals shared and distinct immune responses and metabolism during SARS-CoV-2 and HIV-1 infections

**DOI:** 10.1101/2022.01.10.475725

**Authors:** Tony Pan, Guoshuai Cao, Erting Tang, Yu Zhao, Pablo Penaloza-MacMaster, Yun Fang, Jun Huang

**Author notes:** These authors contributed equally: Tony Pan and Guoshuai Cao.

## Abstract

SARS-CoV-2 and HIV-1 are RNA viruses that have killed millions of people worldwide. Understanding the similarities and differences between these two infections is critical for understanding disease progression and for developing effective vaccines and therapies, particularly for 38 million HIV-1^+^ individuals who are vulnerable to SARS-CoV-2 co-infection. Here, we utilized single-cell transcriptomics to perform a systematic comparison of 94,442 PBMCs from 7 COVID-19 and 9 HIV-1^+^ patients in an integrated immune atlas, in which 27 different cell types were identified using an accurate consensus single-cell annotation method. While immune cells in both cohorts show shared inflammation and disrupted mitochondrial function, COVID-19 patients exhibit stronger humoral immunity, broader IFN-I signaling, elevated Rho GTPase and mTOR pathway activities, and downregulated mitophagy. Our results elucidate transcriptional signatures associated with COVID-19 and HIV-1 that may reveal insights into fundamental disease biology and potential therapeutic targets to treat these viral infections.

**Highlights:**

- COVID-19 and HIV-1^+^ patients show disease-specific inflammatory immune signatures
- COVID-19 patients show more productive humoral responses than HIV-1^+^ patients
- SARS-CoV-2 elicits more enriched IFN-I signaling relative to HIV-I
- Divergent, impaired metabolic programs distinguish SARS-CoV-2 and HIV-1 infections

## Introduction

SARS-CoV-2 and HIV-1 have both claimed millions of lives. SARS-CoV-2 has infected over 290 million people worldwide, resulting in more than 5.4 million deaths by December 2021. (Dong et al., 2020). There are currently over 38 million people living with HIV-1 (PLWH) and over 36 million AIDS-related deaths since the beginning of the AIDS epidemic (UNAIDS, 2021). SARS-CoV-2 and HIV-1 are both RNA viruses and thus exhibit high mutation rates relative to DNA viruses. SARS-CoV-2 and HIV-1 are both highly virulent, but disease progression with these viruses differs substantially. For instance, most of the mortality and morbidity observed with SARS-CoV-2 infection occurs within days of infection, compared to months or years with HIV-1 infection. Furthermore, neutralizing antibody responses are rapidly generated following SARS-CoV-2 infection, but these take many years to develop in PLWH (Cotugno et al., 2021; Dangi et al., 2021b; Stamatatos et al., 2009). Various antiviral treatments have been developed for HIV-1, while treatments for SARS-CoV-2 are still limited. These clinical and immunological differences are driven in part by how the host responds to these distinct viral infections. In the current COVID-19 pandemic, PLWH often have compromised immune systems, which may render them vulnerable to SARS-CoV-2 infection and exhibit suboptimal responses to SARS-CoV-2 vaccination. Thus, a comprehensive immune profiling of SARS-CoV-2 and HIV-1 infections would help us understand the mechanisms by which these two viruses cause diseases and deaths, guiding the discovery of novel therapeutics.

Patients with severe COVID-19 infection typically exhibit “cytokine storms” that are linked to more severe disease outcomes (Ragab et al, 2020). In particular, patients with severe COVID-19 produce high levels of inflammatory cytokines and chemokines including IL-6, IL-10, TNF-α, IFN-γ, and IP-10 (Ragab et al, 2020). Such cytokines are also expressed during the acute response to HIV-1 infection, but a dysregulated cytokine response can persist if HIV-1 infection is left untreated (Deeks et al., 2013). Various immune cell subsets have been also implicated in driving inflammatory responses during HIV-1 infection, namely macrophages and monocytes (Campbell et al., 2014), and their inflammatory roles have also been characterized in COVID-19 (Lee et al., 2020; Liu et al., 2021; Melms et al., 2021). However, the distribution and cell type-specific functions of different immune cell populations (T cells, B cells, natural killer cells, dendritic cells, monocytes) are known to vary across different diseases, conditions, and stages of disease progression (Delorey et al., 2021; MacParland et al., 2018; Travaglini et al., 2020). Thus, studies to specifically compare immune cell populations during HIV-1 and COVID-19 infections at the single-cell level are still lacking.

Given the complexity of the immune system, single-cell RNA sequencing (scRNA-seq) has been extensively deployed to understand the heterogeneity within immune cell subsets (Chow et al., 2021; Gawad et al., 2016; Reyfman et al., 2019; Tang et al., 2009; Treutlein et al., 2014). scRNA-seq has been proven more powerful than most bulk methods in revealing complex cell-cell interactions, regulatory modules, and subpopulation dynamics with the single-cell resolution (Chen et al., 2019a; Goveia et al., 2020; Haque et al., 2017; Kuksin et al., 2021). A main advantage of scRNA-seq is accurate annotation of individual cells. However, most studies utilize manual supervision, which can be subjective and difficult to compare across studies (Liao et al., 2020; Wen et al., 2020). While multiple scRNA-seq atlases have been established on COVID-19, they differ significantly in their granularity and markers used for annotation (Melms et al., 2021; Lee et al., 2020; Wen et al., 2020; Liao et al., 2020). There have been few HIV-1 scRNA-seq profiling studies, and it is unclear how these compare to COVID-19. In addition, different sequencing methods and analysis pipelines render a comparison between HIV-1 and COVID-19 difficult. Therefore, a reliable and accurate integration strategy is needed to transfer and synthesize our understandings from COVID-19 to HIV-1 and generate insights that could lead to better and synergistic treatment of both diseases, which is particularly relevant to HIV-1^+^ patients in the current COVID-19 pandemic.

To date, a study comparing gene expression at the single-cell level following SARS-CoV-2 and HIV-1 infections to better understand the host’s response to infection with these viruses has not been performed. Here, we present a comprehensive strategy to integrate scRNA-seq data of 115,272 single PBMCs from 7 COVID-19 (Wilk et al., 2020), 9 HIV-1^+^ (Kazer et al., 2020; Wang et al., 2020) and 3 healthy patients (10xGenomics, 2020). Our strategy combined the advantages of manual annotation, correlation-based label transfer and deep-learning-based classification to generate a high-quality unified cellular atlas of the immune landscape. Based on this atlas, we compared in detail the phenotypic features and regulatory pathways in each of the major immune compartments (T cells, B cells, natural killer cells, dendritic cells, and monocytes).Overall, our single-cell annotation method provides a straightforward scRNA-seq integration strategy that can be extended in multiple settings beyond the current study. In addition to finding common signatures of inflammation and disrupted mitochondrial function in both COVID-19 and HIV-1, we also found important differences in cell signaling, antibody diversity, IFN-I signaling, and metabolic function. Our findings provide an important resource to better understand the pathophysiological differences between COVID-19 and HIV-1, which may lead to novel molecular targets for the treatment of these diseases.

## Results

### Consensus clustering approach corrects cell type labels and reveals additional cell subsets

To examine the differential antiviral response, we aggregated publicly available scRNA-seq data from PBMCs derived from 7 severe COVID-19 (22,078 cells) (Wilk et al., 2020), 9 HIV-1^+^ (72,364 cells) (Kazer et al., 2020; Wang et al., 2020), and 3 healthy patients (20,830 cells) (10xGenomics, 2020) (Figure 1A, left). In order to identify the cell type-specific responses in each disease, we designed an integrated annotation strategy to improve and synthesize existing cell labels in the respective studies.

**Figure 1:**
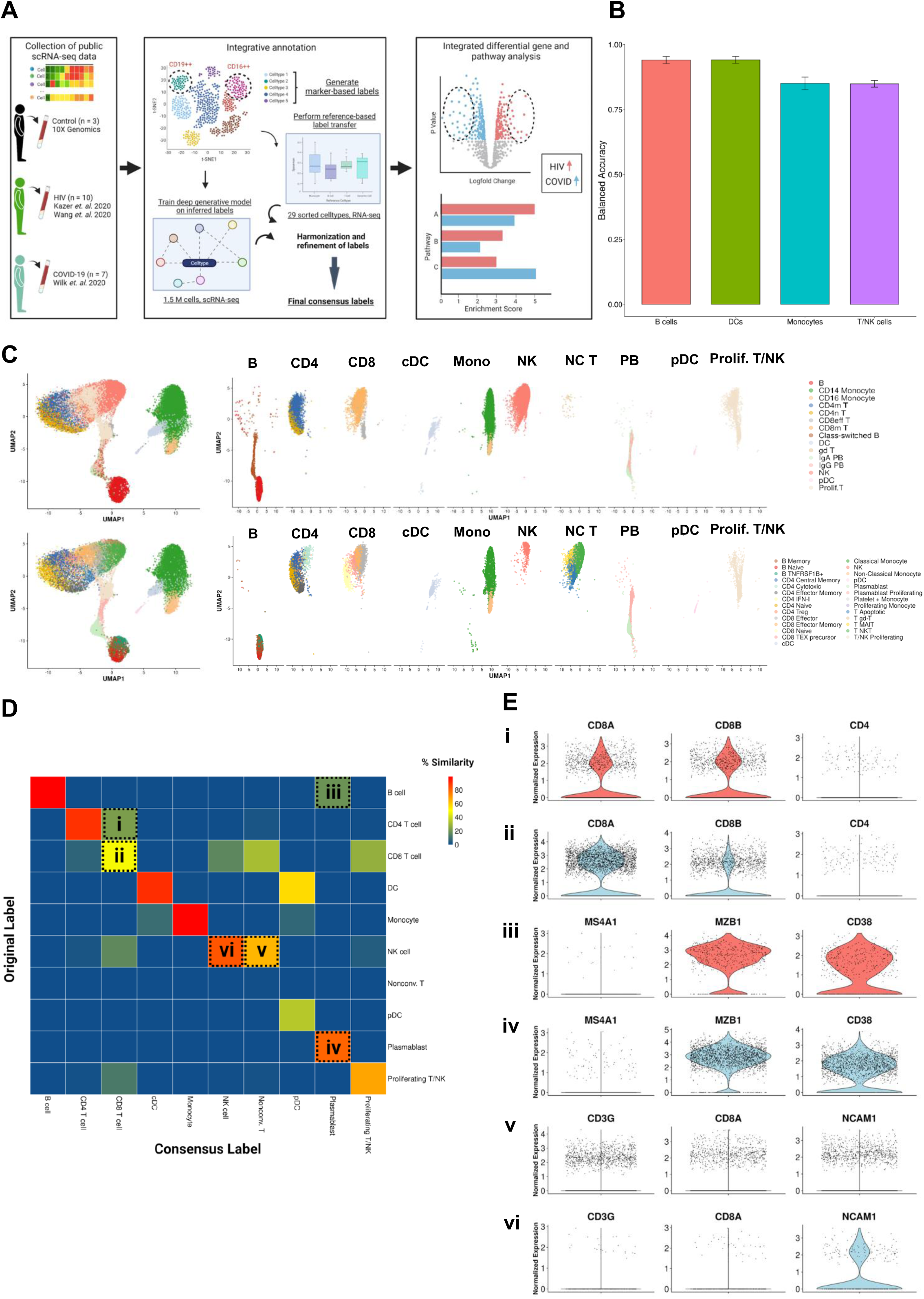
Consensus clustering method to annotate single cell transcriptomic data from multiple sources. (A) Illustrated workflow of data collection, consensus annotation, and downstream analysis. (B) Balanced accuracy of trained scANVI model on cell labels derived from Ren Cell 2021. Error bars denote variation of accuracy across labels within major cell categories. (C) Left: Uniform manifold approximation and projection (UMAP) embeddings of the integrated datasets colored by original (top) and consensus labels (bottom). Right: UMAP embeddings split by major cell categories colored by original (top) and consensus labels (bottom) illustrating the contrast in cell proportions using consensus method. (D) Confusion matrix illustrating percentage overlap of original labels and consensus labels across major cell categories. Percentage overlap was calculated by dividing each cell count by the total number of cells in each column. (E) Violin plots of canonical normalized gene expression of designated cell populations indicated in 1D (rows).

Our integration strategy is based on the combination of three different annotation approaches, namely manual annotation, correlation-based label transfer and deep-learning-based classification (Aran et al., 2019; Xu et al., 2021). Manual annotation based on expression of known biological markers is widely used. However, this method can vary in accuracy because of its subjectivity and varying knowledge of markers (Abdelaal et al., 2019; Pasquini et al., 2021). Correlation-based methods such as SingleR (Aran *et al*., 2019) offer improved accuracy by correlating query scRNA-seq gene expression with bulk RNA-seq data of pure cell populations from healthy donors to transfer known labels but lack the context-specific or disease-specific features. Deep-learning based methods such as scANVI (Xu et al. 2021) train a classifier on context-specific or disease-specific atlas data to generate a probabilistic model that can classify new data independent of technical variation between datasets, making it both highly specific and scalable. However, overfitting can happen if such deep-learning models are left unchecked against known biology. To leverage the advantages of each approach, we performed cell annotation using each of the three methods independently, and then integrated the three sets of labels to produce one final set of consensus labels (Figure 1A, center; see Materials and Methods). Our deep learning annotation resulted in high accuracy across cell types (Figure 1B).

Our integration strategy resulted in 27 total cell types, consisting of 5 B cell subsets, 2 dendritic cell (DC) subsets, 4 monocyte subsets, 7 CD4^+^ T cell subsets, 8 CD8^+^ T cell subsets, and 1 natural killer (NK) cell subset (Figure 1C, bottom). This is a substantial improvement to the 15 cell types provided from the source publication (Wilk et al, 2020), which comprised of 4 B cell subsets, 2 DC subsets, 2 monocyte subsets, 2 CD4^+^ T cell subsets, 4 CD8^+^ T cell subsets, and 1 NK cell subset (Figure 1C, top). Notably, we found greater granularity amongst CD4^+^ T cells, CD8^+^ T cells, and unconventional T cells; we were able to identify unclassified populations: effector memory CD4^+^ T cells, cytotoxic CD4^+^ T cells, IFN-I^+^ CD4^+^ T cells, regulatory CD4^+^ T cells (Tregs), naïve CD8^+^ T cells, precursor exhausted CD8^+^ T cells, NKT cells, MAIT cells, and apoptotic T cells. While the bulk of our consensus cell assignments agreed with the original labels, we discovered critical discrepancies in some cell types. We compared the original label of each cell with their consensus label and summarized the results in a confusion matrix (Figure 1D). This allowed us to identify the specific subpopulations with disagreeing labels and resolve them using canonical gene expression. In the T cell compartment, a significant proportion of CD8^+^ T cells were originally classified as CD4^+^ (Figure 1D ‘i, ii’). When comparing the expression of canonical genes *CD8A*, *CD8B*, and *CD4* in this population (Figure 1E ‘i’) to the expression of the main cluster of CD8^+^ T cells (Figure 1E ‘ii’), we saw that levels were markedly similar, leading us to conclude that they are indeed CD8^+^ T cells. Similarly, we used expressions of *MZ4A1* (a canonical B cell marker), *MZB1*, and *CD38* (canonical plasmablast markers, Figure 1E ‘iii, iv’) to confirm that the population indicated in Figure 1D ‘iii’ are plasmablasts instead of B cells, and the expression of *CD3G*, *CD8A*, and *NCAM1* (a canonical NK cell marker, Figure 1E ‘v, vi’) to confirm that the population indicated in Figure 1D ‘v’ are unconventional T cells instead of NK cells. Our labels also consistently displayed high cluster purity according to their ROGUE score (Liu et al., 2020) (Figure S1D). Overall, our consensus clustering approach allowed us to generate high-resolution labels with improved biological accuracy.

### Integrated landscape of PBMCs from COVID-19, HIV-1^+^ and healthy patients

To compare the full immune landscape across our three conditions, we integrated single-cell data of all 19 patients (7 COVID-19, 9 HIV-1^+^ and 3 healthy patients) into a single UMAP and grouped our 27 consensus clusters into 10 major cell types (Figure 2A). The resulting balanced distribution of cells across the three disease conditions demonstrated the successful integration (Figure 2B upper). The structure of the UMAP reveals four major cell populations. The top left cluster comprises CD4^+^ and CD8^+^ T cells, innate-like T cells, NK cells, and proliferating cells; the central and bottom clusters comprise primarily plasmablasts and B cells; and the rightmost cluster consists of monocytes and dendritic cells (Figure 2B lower). Because scRNA-seq results can vary significantly due to confounding factors across batches, we explicitly regressed out any patient specific effects, resulting in a representative distribution of every cell type across each patient (Figure 2C). When comparing the frequency of major cell types across conditions, we found that COVID-19 and HIV-1^+^ patients had elevated counts of innate-like T cells, CD8^+^ T cells, and monocytes compared to healthy controls (Figure 2D), demonstrating the recruitment of inflammatory cells with either viral infection.

**Figure 2:**
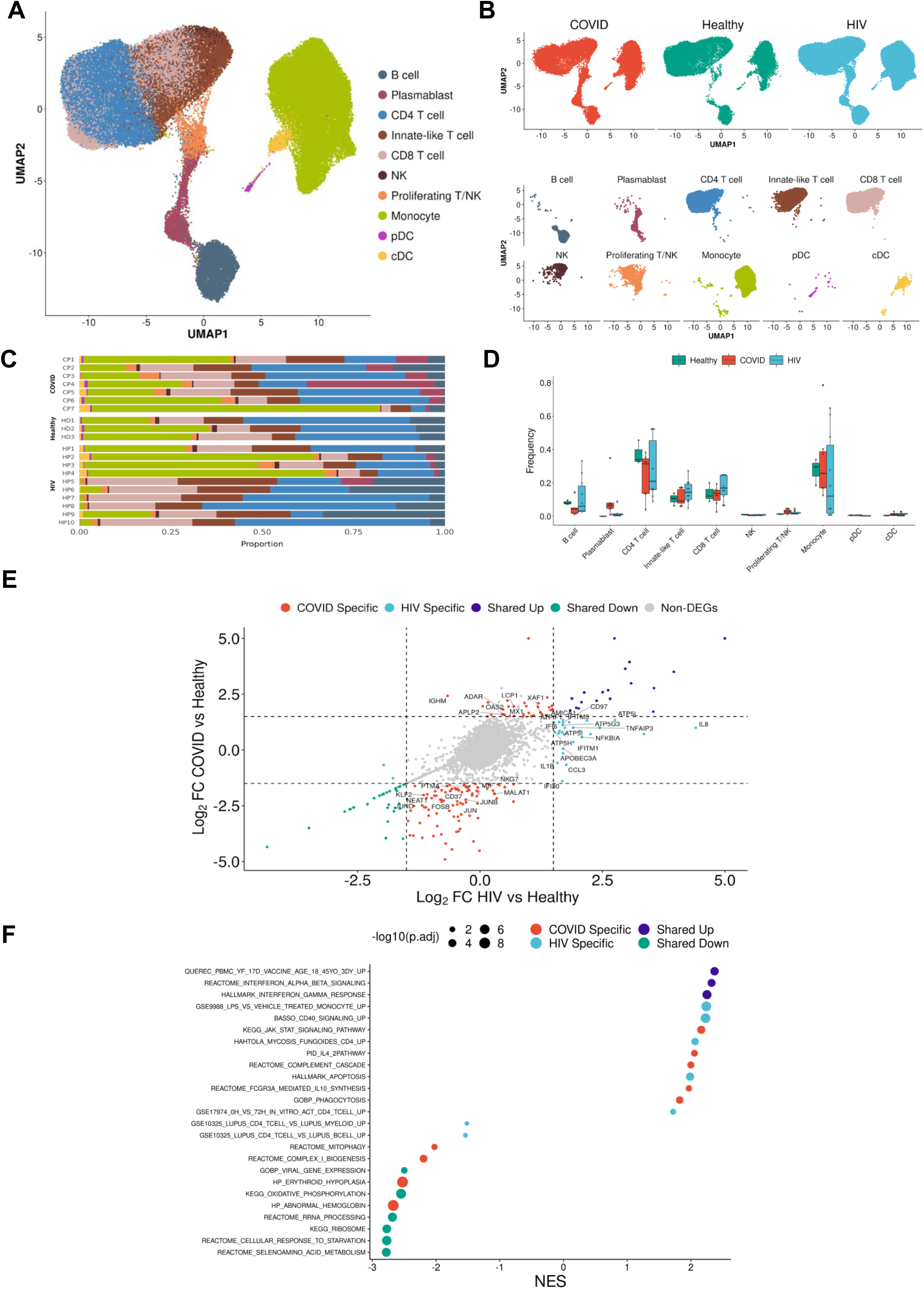
Integrated single-cell landscape of PBMC in HIV, COVID-19, and healthy controls. (A) UMAP embeddings of integrated HIV-1^+^ and COVID-19 patients together with healthy controls colored by major cell populations. (B) Top: UMAP split across disease conditions after regressing out patient-specific effects using the Harmony algorithm. Bottom: UMAP highlighting distribution of major cell populations. (C) Stacked bar plots of the relative frequency of major cell populations present in each patient. CP: COVID-19 patient. HP: HIV-1 patient. HD: Healthy donor. (D) Box plots of the relative frequency of major cell populations present in each condition. Each point represents one patient. (E) Double differential gene expression plot of genes that are differentially expressed between COVID-19 patients compared to healthy controls (Y-axis) or differentially expressed between HIV-1^+^ patients compared to healthy controls (X-axis). Log2FC: Log-2-fold change. (F) Dot plot of enriched biological pathways from differentially expressed genes that were found to be upregulated (right, positive) or downregulated (left, negative) compared to healthy controls. Size of dot corresponds to adjusted P value of enriched pathway. NES: Normalized enrichment score.

To further contrast COVID-19 and HIV-1^+^ patients, we performed differential gene expression analysis to derive two sets of differentially expressed genes (DEGs): one set comparing gene expression on COVID-19 versus healthy controls, and the other comparing HIV-1 versus healthy controls. We compiled all DEGs with an adjusted p-value < 0.05 and compared their differential expression in COVID-19 versus healthy controls and HIV-1 versus healthy controls (Figure 2E). We also identified two distinct sets of disease-specific DEGs. HIV-1^+^ patients exhibited substantial upregulation of *IL8*, *CCL3*, and *NFKBIA*, which have been implicated in the antiviral response and inflammation. In contrast, patients with COVID-19 showed upregulation of *OAS2*, *XAF1*, and *MX1*, which are part of the type-I interferon (IFN-I) signaling pathway (Figure S2A). OAS proteins function to degrade double-stranded RNAs which are intermediates during coronavirus replications (Choi et al., 2015). A recent genome-wide association study (GWAS) reported a significant association between genetic variants in human OAS genes and COVID-19 severity (Pairo-Castineira et al., 2021). COVID-19 patients downregulated genes involved in the AP-1 transcription factor pathway (including *JUN*, *JUNB*, *JUND*, and *FOSB)* as well as HLA genes (including *HLA-E*, *HLA-DRB1*, *HLA-DRA*, and *HLA-DPB1*) (Figure S2C). We found a joint downregulation of *LTB* (which encodes for lymphotoxin-B, an inflammatory protein that plays a role in lymphoid tissue development) (Lu and Browning, 2014) and *KLF2* (which regulates the differentiation and function of immune cells) (Jha and Das, 2017). We also found a joint upregulation of interferon-associated genes including *ISG15*, *IFI27,* and *IFITM3* (Figure S2B).

To further explore these genes, we performed gene set enrichment analysis (GSEA) on each set of the DEGs (Figure 2F). We found that both COVID-19 and HIV-1^+^ patients were highly enriched in interferon-alpha/beta (IFN-α/β) signaling and interferon-gamma (IFN-γ) signaling, despite a lack of shared upregulated genes. We also found a shared downregulation of ribosome and oxidative phosphorylation (OXPHOS) pathways, which indicates that both infections cause a shift in metabolic function, potentially due to viral hijacking of cellular metabolic machinery or host responses. Moreover, we identified disease-specific pathways enriched only in either disease. Most notably, cells from COVID-19 patients were enriched in JAK-STAT signaling, IL-4 signaling, and IL-10 production, which are known to play an important role in the antiviral response (de la Rica et al., 2020; Lu et al., 2011; Satarker et al., 2021). In contrast, cells from HIV-1^+^ patients were enriched in CD40 signaling and CD4^+^ T cell activation, as well as apoptosis. In conclusion, while our analysis of the immune landscape in COVID-19 and HIV-1 revealed similarities in the biological processes that regulate each disease, expression of the specific genes involved are very different, revealing nuanced differences in how immune cells are responding to these two viral infections.

### Inflammatory innate immune cells are a hallmark of both COVID-19 and HIV-1 infections

Innate immune cells play a vital role in the response to viral infection. Monocytes and DCs are capable of directly sensing viral particles via pattern-recognition receptors, triggering intracellular signaling events to initiate a cytokine and chemokine-mediated inflammatory response (Takeuchi and Akira, 2007). Additionally, such innate cells modulate the adaptive response to viral infection through cell-cell interactions or soluble factors, which help polarize T cells toward a Th1 phenotype (Gasteiger and Rudensky, 2014). These functions have been shown to be critical in mounting immune responses in both COVID-19 and HIV-1 (Campbell *et al*., 2014; Coleman and Wu, 2009; Kasuga et al., 2021; Schultze and Aschenbrenner, 2021). While a strong inflammatory monocyte response is a hallmark of both diseases, important differences remain to be elucidated. For example, while peripheral monocytes have been shown to be susceptible to HIV-1 infection, little is known for SARS-CoV-2 infection (Kedzierska and Crowe, 2002; Zhang et al., 2021). Additionally, prior studies have identified polyfunctional monocytes that have been associated with elite response in HIV-1 (Kazer et al, 2020), and it is unclear whether these populations are conserved across viral infections or specific to HIV-1. Thus, we sought to closely examine the transcriptomic differences between DCs and monocytes in COVID-19 and HIV-1^+^ patients.

We subsetted out only the DCs and monocytes and performed integration and clustering. We identified 6 total clusters (Figure 3A): conventional dendritic cells (cDCs, *CD1C*^high^ *FCER1A*^high^ *CLEC10A*^high^), plasmacytoid dendritic cells (pDCs, *CLEC4C*^high^ *IL3RA*^high^ *TCF4*^high^), classical monocytes (*CD14*^high^ *FCGR3A*^low^), nonclassical monocytes (*CD14*^low^ *FCGR3A*^high^), proliferating monocytes (*CD14*^high^ *MKI67*^high^ *TOP2A*^high^), and a mixed platelet and monocyte cluster (*PPBP*^high^ *PF4*^high^ *CD14*^high^) (Figure 3B). We found that HIV-1^+^ patients exhibited significantly higher proportions of cDCs compared to COVID-19 patients (Figures 3C and 3D), which could be explained by the dual role of cDCs in HIV-1 infection: in addition to presenting HIV-1 antigens, cDCs also play an antiviral role (Manches et al., 2014). We also found that HIV-1^+^ patients exhibited significantly higher frequencies of nonclassical monocytes (Figures 3C and 3D), which have been shown to massively expand in the peripheral blood in response to immune activation by HIV-1 infection (Campbell et al., 2014). Differential gene expression analysis revealed a shared inflammatory phenotype across both HIV-1 and COVID-19 sharing genes such as *IFITM3* and *IFI27*, which play a role in driving IFN-I signaling (Figures 3E, 3F, and S3C). However, we once again found the majority of the contributing genes to be virus-specific. Monocytes of HIV-1^+^ patients highly express genes associated with proinflammatory cytokines including *IL8*, *IL1B,* and *CCL3*, all of which play a role in the acute viral response and immune cell recruitment (Figure 3E and S3A). Monocytes of COVID-19 patients highly express genes associated with inflammation including *IL17RA* and *MX2* in addition to JAK-STAT associated genes *STAT2* and *STAT6* (Figure S3B). Interestingly, while cDCs in HIV-1 patients did slightly upregulate *SAMHD1*, an antiretroviral protein shown to be effective in inhibiting early HIV-1 infection, it was much more highly upregulated by cDCs in COVID-19 patients. In agreement with previous studies, GSEA analysis revealed joint upregulation of inflammatory pathways such as IFN-I response and IFN-α/β signaling (Figure 3F). However, we also found a greater diversity of inflammatory response in COVID-19 compared to HIV-1. Only COVID-19 monocytes and DCs upregulated signaling by IL-20, IL-2, IL-6, KIT, and JAK-STAT, suggesting that innate immune cells may be much more active and cytotoxic in COVID-19 compared to in HIV-1 (Figure 3F). Several of these cytokines, including IL-6, have been found to be overexpressed in COVID-19 patients and shown to be positively correlated to disease severity (Costela-Ruiz *et al*., 2020; Jones and Hunter, 2021; Ma et al., 2021; Rubin et al., 2021; Weisberg *et al*., 2020).

**Figure 3:**
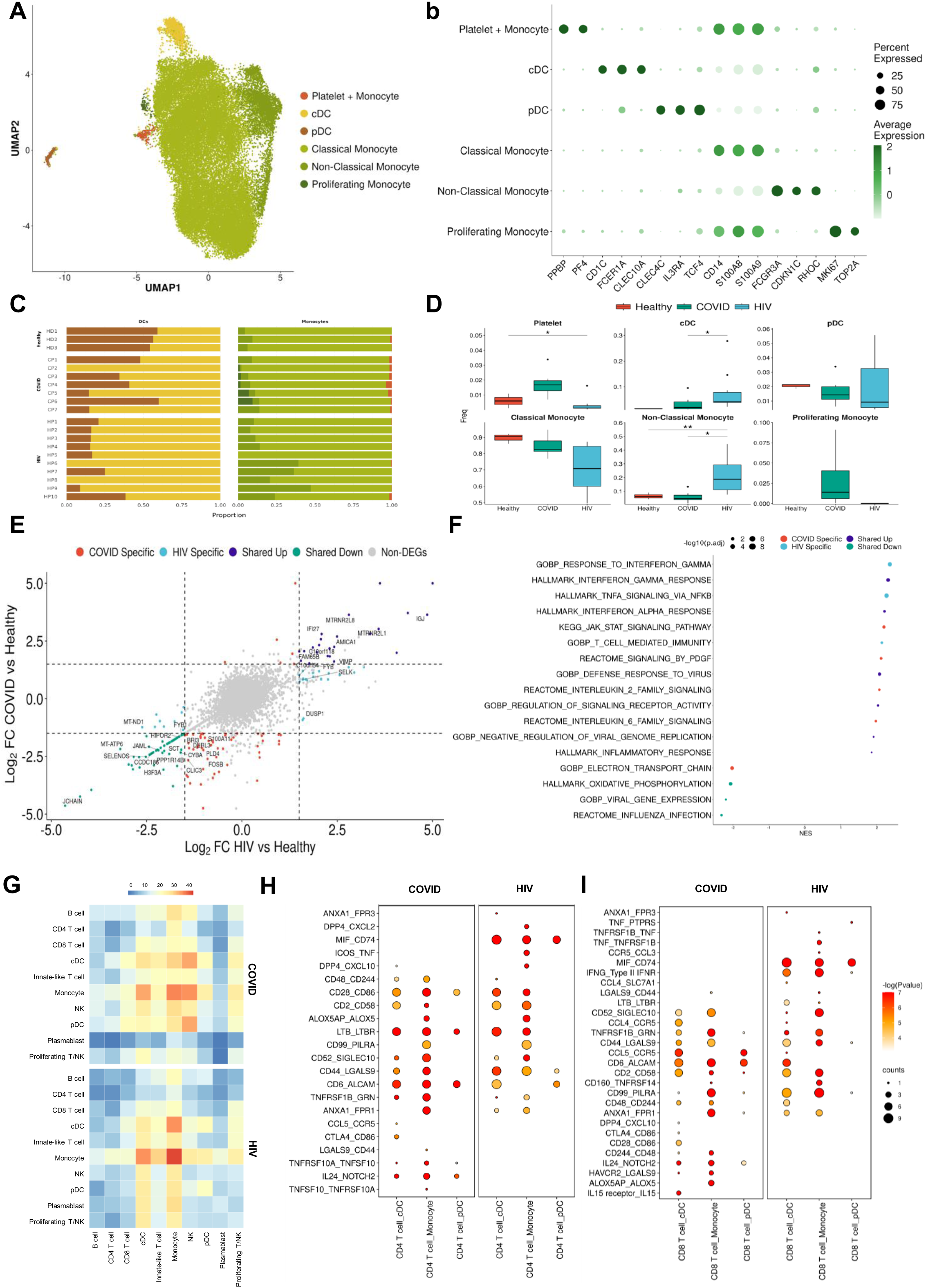
Monocytes in COVID-19 and HIV-1 share inflammatory signatures. (A) UMAP embeddings of monocytes and DCs colored by subtype. (B) Dot plot of canonical monocyte and DC marker expression across subtypes. (C) Stacked bar plots of the relative frequency of subtypes present in each patient. (D) Box plots of the relative frequency of subtypes present in each condition. Significance testing was doing using student’s t-test. “*”: P < .05, “**”: P < .01, (E) Double differential gene expression plot of genes that are differentially expressed between COVID-19 patients compared to healthy controls or differentially expressed between HIV-1^+^ patients compared to healthy controls. (F) Dot plot of enriched biological pathways from differentially expressed genes that were found to be upregulated (right, positive) or downregulated (left, negative) compared to healthy controls. (G) Heatmap of the number of receptor-ligand interactions between each cell type in COVID-19 patients (top) and HIV-1^+^ patients (bottom). (H) Dot plot of selected receptor-ligand interactions between CD4^+^ T cells and monocytes/DCs in COVID-19 patients (left) versus HIV-1^+^ patients (right). Color of each dot corresponds to the inverse log of the P value of the interaction. Size of the dot corresponds to the number of patients the interaction was found to be significant in. (I) Dot plot of selected receptor-ligand interactions between CD8^+^ T cells and monocytes/DCs in COVID-19 patients (left) versus HIV-1^+^ patients (right). Color of each dot corresponds to the inverse log of the P value of the interaction. Size of the dot corresponds to the number of patients the interaction was found to be significant in.

Given these differences in the functional profiles of DCs and monocytes across the two viral infections, and the frequent interactions with the adaptive immune system as antigen presenting cells, we surmised that they may also play a divergent role in mediating the adaptive immune response. To investigate this interaction network, we utilized CellphoneDB (Efremova et al., 2020) to determine the putative receptor-ligand interactions based on their gene co-expression patterns on pairs of cell types (Figure 3G, Figures S5A-D). We found significant differences in highly-interacting pairs in HIV-1 compared to COVID-19. We found that despite having a lower relative frequency in COVID-19 compared to HIV-1, cDCs in COVID-19 patients had more frequent interactions with monocytes, NK cells, and T cells but less frequent interactions with plasmablasts compared to cDCs in HIV-1^+^ patients. While monocytes had a high frequency of interactions in both COVID-19 and HIV-1^+^ patients, we found that DC-T cell and monocyte-T cell interactions were noticeably enriched in COVID-19 (Figures 3G, S6A, and S6B). To investigate this further, we selected the most significant CD4^+^ and CD8^+^ T cell interactions with monocytes/DCs across the two infections (Figures 3H, 3I, and S7A-H). We found a large number of costimulatory and inflammatory interactions shared across cell types and diseases, notably *CD28*-*CD86* (which provides a critical costimulatory signal for T cells) (Hui et al., 2017), *CD6*-*ALCAM* (which drives immune synapse formation and activation, and migration in CD4^+^ T cells) (Ampudia et al., 2020), and *TNFRSF1B*-*GRN* (which drives apoptosis and inflammation) (Ward-Kavanagh et al., 2016). We also found enrichment of the inhibitory interaction *CD99*-*PILRA* (which curbs NK-like cytotoxicity). Migratory inhibitory factor (MIF), which is responsible for the pathogenesis of viral infection, is known to be present in higher concentration in the periphery of HIV-1^+^ patients (Regis et al., 2010). We found the *MIF*-*CD74* interaction between cDCs/monocytes and CD4^+^/CD8^+^ T cells to be unique and highly prevalent across HIV-1^+^ patients, but not COVID-19 patients (Figures 3H and 3I). We also found the interferon IFNγ-IFNγ receptor interaction to be uniquely upregulated in HIV-1 CD8^+^ T cells. In COVID-19, we found the inflammatory *NOTCH2*-*IL24* interaction (which induces *STAT1* and *STAT3* to regulate cell proliferation and survival (Ouyang and O’Garra, 2019) and the inhibitory interactions *CTLA4*-*CD86* and *HAVCR2*-*LGALS9*. Altogether, this analysis suggests that innate-induced inflammation is present in both diseases but are likely driven by very different genes and cell-cell interactions.

### COVID-19 exhibits a stronger plasmablast and antibody response compared to HIV-1

B cells are the primary effectors of the humoral antiviral immune response (Upasani et al., 2021). To investigate if B cells from COVID-19 and HIV-1^+^ patients exhibited distinct transcriptional signatures, we performed integration and clustering on B cell and plasmablast populations, identified by overexpression of *CD19*/*MS4A1* and *CD38* respectively (Figure 4A). We found 5 total subpopulations: naïve B cells (*TCL1A*^high^ *IGHD*^high^ *CD27*^low^), memory B cells (*TCLA1*^low^ *CD27*^high^ *AIM2*^high^), *TNFRSF1B*^+^ B cells (*TNFRSF1B*^high^ *CD84*^high^), plasmablasts (*CD38*^high^*XBP*1^high^), and proliferating plasmablasts (*CD38*^high^ *MKI67*^high^ *TOP2A*^high^) (Figure 4B). Consistent with prior COVID-19 studies that have shown extensive plasmablast expansion in patients (Bernardes et al., 2020; De Biasi et al., 2020; Kuri-Cervantes et al., 2020), we found that COVID-19 patients have significantly higher proportions of plasmablasts and proliferating plasmablasts compared to healthy controls (Figures 4C and 4D). Consistent with this result, we found that COVID-19 patients have significantly lower proportions of naïve and memory B cells compared to healthy controls (Figures 4C and 4D). Interestingly, we found a significant enrichment of *TNFRSF1B*^+^ B cells in HIV-1^+^ patients (Figure 4D). These B cells are a subset of effector memory-like B cells given their expression memory B-cell marker genes (intermediate expression of *AIM2* and *CD27*) and upregulation of *TNFRSF1B* and *CD84*. CD84 has been shown to be upregulated on a subset of memory B cells that exhibit higher levels of proliferation and has been associated with B cell activation and signal transduction (Tangye et al., 2002), while *TNFRSF1B* encodes for a TNF-receptor protein that has been known to induce TNF-mediated apoptosis. While HIV-1^+^ patients also had increased proportions of plasmablast and proliferating plasmablast subsets compared to healthy controls, these responses were more moderate compared to COVID-19 patients. We found COVID-19 plasmablasts to express higher levels of *MKI67*, *IGHM*, *CCR2*, and *XBP1* compared to HIV-1 plasmablasts, suggesting their elevated proliferation and maturation (Figure S4D).

**Figure 4:**
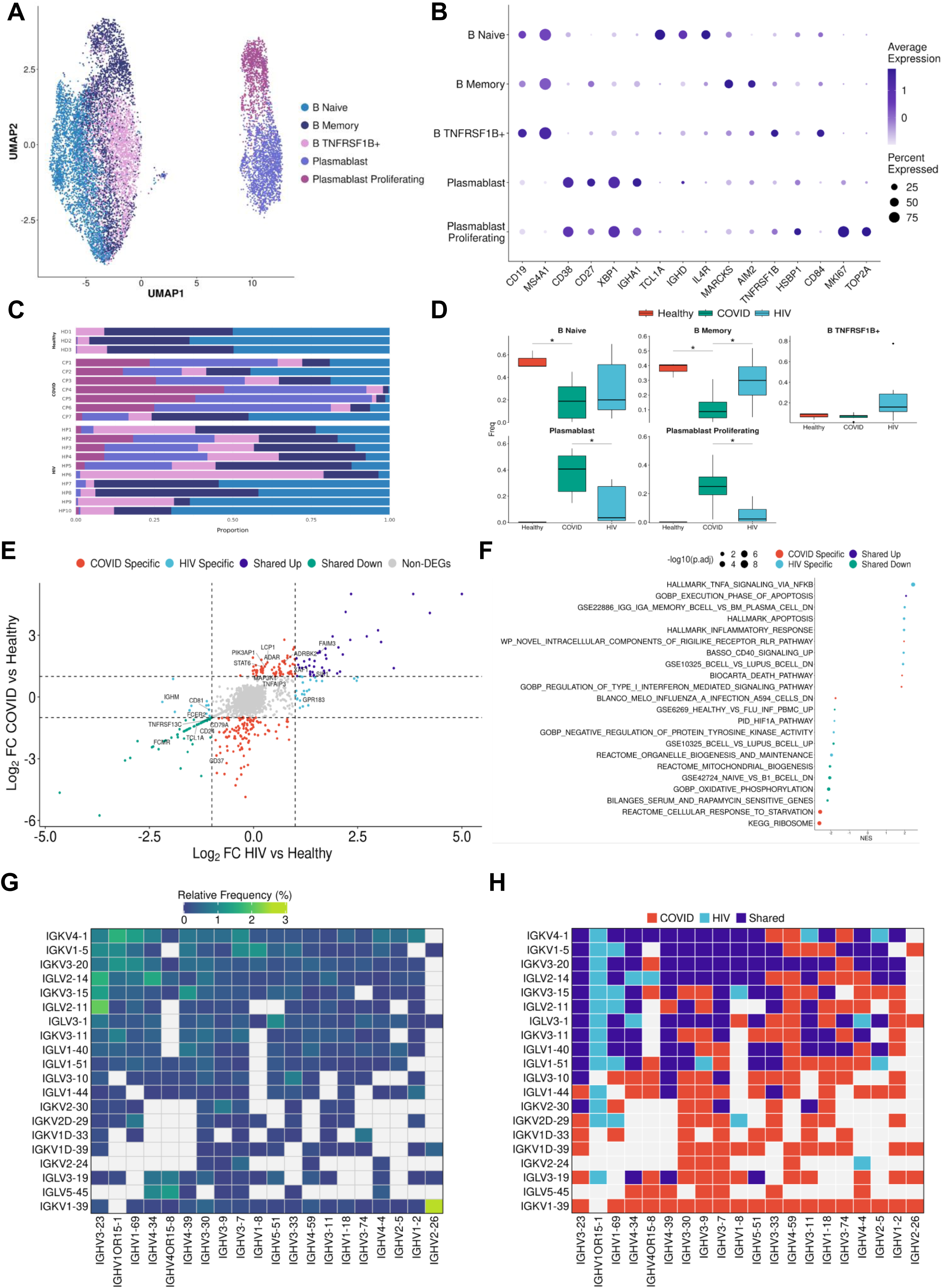
B cells in COVID-19 show more robust plasmablast response and antibody diversity relative to HIV-1. (A) UMAP embeddings of B cells colored by subtype. (B) Dot plot of canonical B cell marker expression across subtypes. (C) Stacked bar plots of the relative frequency of subtypes present in each patient. (D) Box plots of the relative frequency of subtypes present in each condition. Significance testing was doing using student’s t-test. “*”: P < .05. (E) Double differential gene expression plot of genes that are differentially expressed between COVID-19 patients compared to healthy controls or differentially expressed between HIV-1 patients compared to healthy controls. (F) Dot plot of enriched biological pathways from differentially expressed genes that were found to be upregulated (right, positive) or downregulated (left, negative) compared to healthy controls. (G) Heatmap of top 20 light chain (Y axis) and heavy chain (X axis) combinations found in HIV-1^+^ and COVID-19 patients. H) Heatmap indicating the light chain/heavy chain combinations that are either unique to HIV-1 (light blue), COVID-19 (red), or shared across the two diseases (dark blue).

To further compare B cells and plasmablasts between COVID-19 and HIV-1 patients, we performed dual differential gene expression analysis, relative to healthy controls (Figure 3E). We found joint upregulation of SIK1, a gene that regulates cell cycling and plays a role in plasmablast maturation. We also found joint upregulation of genes involved in apoptosis and activation including *TNFAIP3*, *XAF1*, and *LCP1*, as well as *ADAR*, which has been implicated in viral RNA replication (Zhu et al., 2020). Interestingly, we found joint downregulation of several markers that are conventionally expressed on B cells including *CD24*, *CD37*, *CD40*, and *CD79a*, which play key roles in BCR signaling and B cell regulation. We also found downregulation of *FCER2*, *FCMR*, *LTB*, and *TNFRSF13*, which help regulate cell differentiation and maintain cellular homeostasis. Taken together, these genes suggest that B cells in both COVID and HIV-1 are actively responding to viral infection, and as a result they exhibit a drastic shift away from homeostasis.

We also observed enrichment of COVID-19-specific signaling genes such as *MAP3K1* (Figure 4E), which helps activate JNK and ERK pathways, and *STAT6*, which is involved in IL-4 and IL-13 signaling (de la Rica *et al*., 2020; Goel et al., 2021b). We found upregulation of activation markers *TRFC, CD80, CD86, IL10RA,* and *CD40* in HIV-1 B cells (Figure S3E). GSEA analysis reinforced these findings, as we found pathways relevant to apoptosis to be enriched in both COVID-19 and HIV-1 B cells and plasmablasts (Figure 4F). Consistent with our cell proportion analysis, we found terms related to plasmablasts to be positively enriched in both COVID-19 and HIV-1, while terms related to B cells to be negatively enriched. We also found several important pathways specific to HIV-1: CD40 signaling, which regulates the activation of the noncanonical NF-κB and JNK signaling pathways (Hömig-Hölzel et al., 2008); TNF-α signaling via NF-κB, and the inflammatory response, suggesting that NF-κB may play a central role in regulating the humoral immune response in HIV-1.

Given the strong B cell and plasmablast responses in both diseases, we then sought to explore the antibody diversity across the two viral infections. We mapped the top immunoglobulin light chain (IGVL) and immunoglobulin heavy chain (IGVH) combinations present to determine the most frequent IGVL-IGVH pairings. We then calculated the frequency of the top combinations that were found in either COVID-19 or HIV-1 B cells and plasmablasts (Figure 4G). We categorized each combination as disease-specific if at least 1 cell expressed that combination in that given patient and shared if it was found in a patient from both diseases. From the top combinations, 150 were unique to COVID-19, 29 were unique to HIV-1, and 110 combinations were shared, suggesting that the plasmablast response to produce antibodies is not only stronger in COVID-19, but also more diverse (Figure 4H). Out of the top 20 IGVH and IGVL combinations, we found IGKV1-39/IGHV2-26 (a COVID-19 specific combination) to be the most frequent combination, which could offer insight into potential broadly neutralizing antibody (bnAb) design. Previous studies have shown that the BCR diversity was significantly reduced in COVID-19 patients compared to healthy controls in addition to being skewed toward different V gene segments; namely, the CDR3 sequences of heavy chain in clonal BCRs in COVID-19 patients were more convergent than that in healthy controls (Jin et al., 2021). Here we show that the antibody repertoire also differs when comparing COVID-19 to HIV-1 despite both being viral infections. The neutralizing antibody response against HIV-1 has been known to be ineffective due to a variety of factors (Moir and Fauci, 2009). We show that there is a distinctly lower number of HIV-1-unique antibody combinations, which could imply the lack of diversity in the antibody response to HIV-1.

### T cells in COVID-19 and HIV-1 patients exhibit different IFN-I profiles

T cell responses are important for viral control (Buggert et al., 2018; Locci et al., 2013). CD4^+^ T cells are directly targeted by HIV-1 resulting in CD4 T cell depletion. Although SARS-CoV-2 infection occurs mostly in the respiratory tract, CD4^+^ T cell depletion has also been reported in COVID-19 patients (Tan et al., 2020). Although the specific cause of CD4^+^ T cell depletion in COVID-19 has yet to be determined, the excess inflammation induced by cytokines such as TNF-α have been proposed as a trigger for T cell apoptosis (Peng et al., 2020). These similarities and differences in T cells motivated a closer investigation into the specific subpopulations that represent the T cell response in both infections.

After integration and clustering of only T cells and NK cells, we found 16 subpopulations in total (Figures 5A and 5B). Among those, three subpopulations were particularly noteworthy. We found an increase in IFN-I^+^ CD4^+^ cells in both COVID-19 and HIV-1^+^ patients, showing high expression of IFN-I-stimulated genes *ISG15* and *IFIT3*, as well as *IL7R*, which were reported to be upregulated by IFNβ in CD4^+^ T cells (Hoe et al., 2010). The second is apoptotic T cells (Figures 5A and 5B), which feature a high expression of mitochondrial genes characteristic of dying cells (Zhu *et al*., 2020). These two clusters are enriched in both COVID-19 and HIV-1^+^ patients compared to healthy controls, although COVID-19 patients have an even higher proportion of IFN-1^+^ CD4^+^ T cells compared to HIV-1^+^ patients (Figures 5C and 5D). The enrichment of apoptotic T-cell subpopulations in both groups is consistent with the notion that HIV-1 infection leads to low levels of CD4^+^ T cells through pyroptosis of abortively infected T cells (Doitsh et al., 2014) and apoptosis of uninfected bystander cells (Garg et al., 2012) as well as the observation that severe COVID-19 patients frequently experienced lymphopenia (Chen et al., 2020b). Finally, there exists a third cytotoxic CD4^+^ T-cell subpopulation, featuring elevated markers of cytotoxicity (*GZMH*, *GNLY*, *NK7G*, *PRF1*, and *GZMB*), indicating a more cytotoxic phenotype (Figures 5A and 5B). The frequency of cytotoxic CD4^+^ T cells is dramatically increased for both cohorts and is higher in COVID-19 patients, suggesting that viral infection can induce CD4^+^ T cell cytotoxicity and the effect is stronger in SARS-CoV-2 infection (Figures 5C and 5D). The frequency of naïve T cells decreased significantly in COVID-19 and HIV-1^+^ patients; correspondingly, effector memory T cells and effector T cells are enriched in both patients, with more enrichment in HIV-1^+^ patients, suggesting these two viral infections can cause different levels of T-cell activation (Figures 5C and 5D).

**Figure 5:**
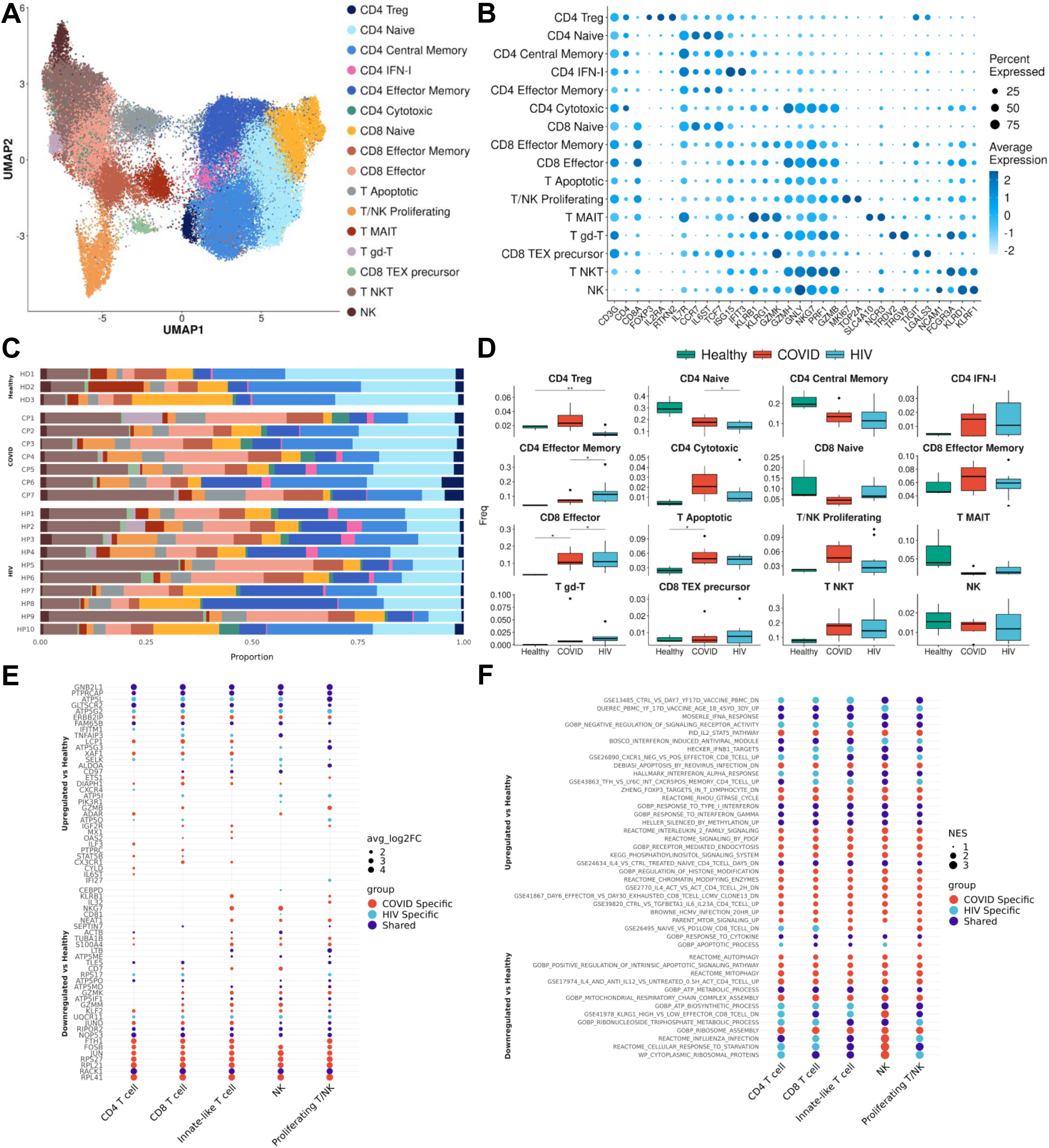
T cells in COVID-19 and HIV-1 show varied IFN-I and activation signatures. (A) UMAP embeddings of T cells colored by subtype. (B) Dot plot of canonical T cell marker expression across subtypes. (C) Stacked bar plots of the relative frequency of subtypes present in each patient. (D) Box plots of the relative frequency of subtypes present in each condition. Significance testing was doing using student’s t-test. “*”: P < .05, “**”: P < .01. (E) Dot plots of the key genes differentially upregulated (top) or downregulated (bottom) compared to healthy controls. (F) Dot plot of enriched biological pathways from differentially expressed genes that were found to be upregulated (top) or downregulated (bottom) compared to healthy controls.

With further gene enrichment analysis, we also found that various T-cell activation-associated genes to be upregulated. *PTPRCAP*, which is associated with the key regulator of T lymphocyte activation *CD45*, as well as *CD97*, which plays an important role in T cell activation, was consistently upregulated in T cells from both COVID-19 and HIV-1^+^ patients. COVID-19 subsets additionally expressed *LCP1*, *STAT5B*, and *ILF3*, which are involved in T-cell activation and signaling. We also found that subsets from both diseases express high levels of genes encoding for inflammatory proteins and chemokines; however, the specific upregulated genes were largely different. COVID-19 subsets expressed high levels of *GZMB* and *CXC3CR1*, suggesting increased cytotoxicity and terminal effector function. HIV-1 subsets upregulated *CXCR4* and *TNFAIP3*, which modulate cell proliferation and initiate inflammatory immune responses, respectively. IFN-stimulated, disease-specific genes are also observed to be upregulated in these two diseases; for example: *XAF1* is specifically upregulated in COVID-19 patients while *IFITM1* is specifically upregulated in HIV-1^+^ patients. Interestingly, While *IFITM1* can inhibit HIV-1 infection by interfering with its replication and entry (Lu *et al*., 2011), *GLTSCR2* is also upregulated, a protein central to viral replication (Wang et al., 2016), indicating the complex interactions between virus and host. In addition to its antiviral capability, *XAF1* can enhance IFN-induced apoptosis. Another interesting finding is the specific down-regulation of *NKG7* and *NEAT1* in innate-like T cells and NKs in COVID-19 patients. Since *NKG7* is important for cytotoxic degranulation and downstream inflammation and *NEAT1* is an activator of the NLRP3 inflammasome, their downregulation may indicate their diminished inflammatory state in COVID-19 patients (Chen et al., 2019b; Malarkannan, 2020) (Figure 5E). Pathway enrichment further revealed enhanced activation and cytokine signaling, namely IFN-α and IFN-γ. We found diverse signaling pathways and general activation pathways associated with COVID-19 subsets, which were enriched in IL2/STAT5, PDGF, and MTOR signaling (Figure 5F). HIV-1 subsets, in contrast, exhibited a less terminally differentiated phenotype, and upregulated IFN-β signaling (Figure 5F).

Altogether, our results revealed a shared activated profile characterized by a robust IFN-I response in T cells from both COVID-19 and HIV-1^+^ patients, relative to healthy donors, though important differences remain between these two viral infections. The stronger apoptosis signature in COVID-19 patients could be driven by elevated IFN-I signaling. These findings motivated further investigation into IFN-I signaling in the two diseases as detailed in following sections.

### IFN-I response is correlated with distinct biological functions in COVID-19 versus HIV-1

Throughout our analysis, we repeatedly found genes and pathways related to IFN-I signaling to be jointly upregulated in both COVID-19 and HIV-1 immune cells compared to healthy controls. This was expected, since IFN-I is produced during viral infections, driving the transcription of IFN-stimulated genes leading to the production of effector molecules capable of curtailing viral infection (McNab et al., 2015). However, we found significant disease-specific differences, as the majority of IFN-I genes were differentially upregulated in one disease or the other (Figure 6A). We categorized these three groups of genes into modules and calculated the module scores across cells in each disease for clearer comparisons (Figure 6B, top). As anticipated, the expression of each module was found to be highest in their corresponding diseases. We found that HIV-1 monocytes and cDCs exhibited the highest associated IFN-I score, suggesting that they may be primarily responsible in driving the IFN-I response (Figure 6B, bottom). We examined the cell type-specific expression of important genes and found that the effector molecule *CCL5* was jointly expressed in CD8^+^ T cells in both diseases. In addition, IFI30 was specifically upregulated in monocytes during HIV-1 infection, whereas *SLAMF7* and *IFIT3* were upregulated in plasmablasts and T cells from COVID-19 patients respectively (Figure 6C).

**Figure 6:**
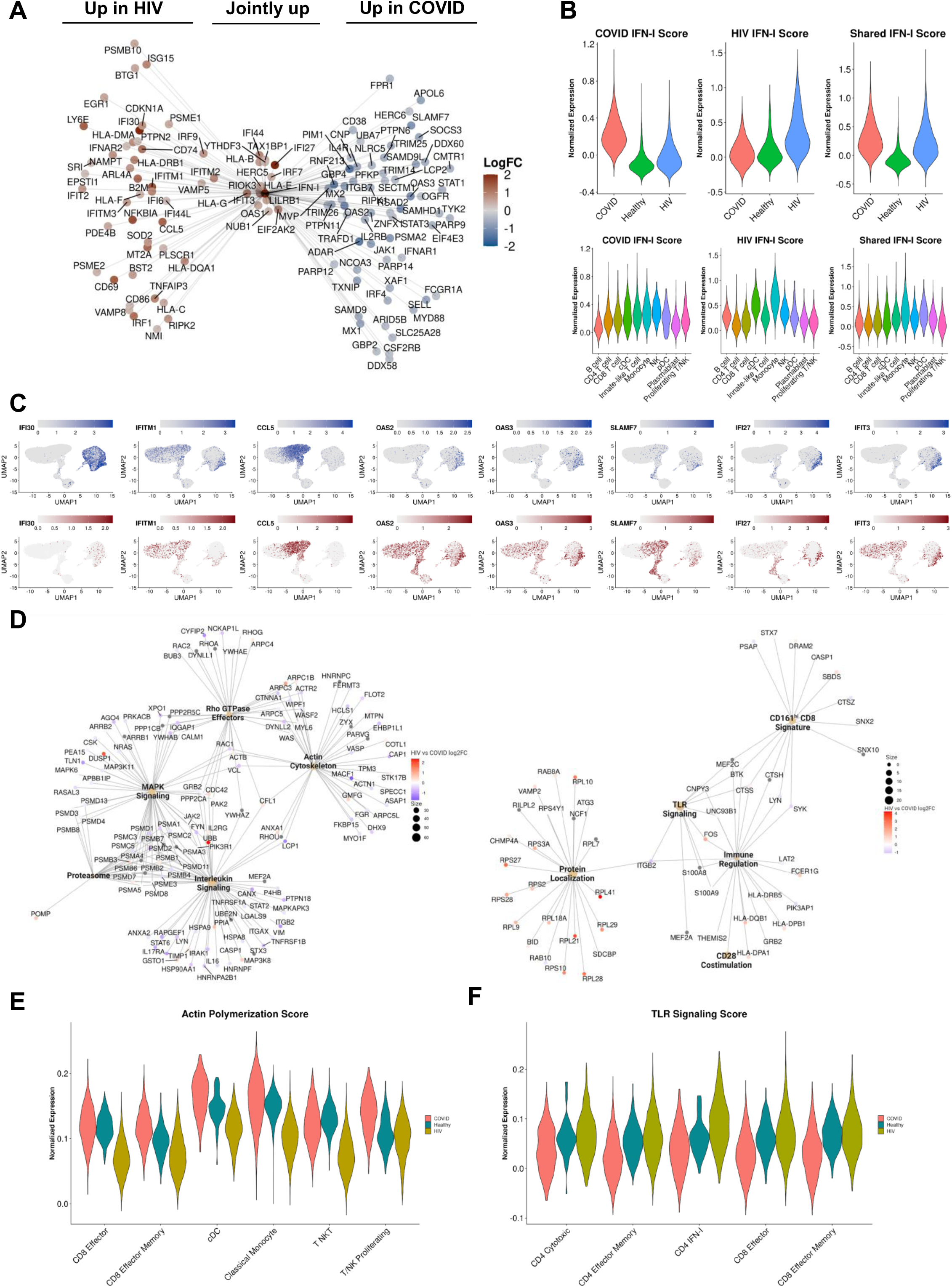
IFN-I signaling is correlated with divergent biological functions in COVID-19 versus HIV-1. (A) Network plot of genes related to IFN-I signaling that are differentially upregulated in COVID-19 (Right), HIV-1 (Left), or jointly upregulated compared to healthy controls. Genes are colored based on Log-2-fold change in expression in HIV-1^+^ versus COVID-19 patients. (B) Violin plots of the normalized gene expression of the three IFN-I signatures in A), split across conditions (top) and major cell populations (bottom). (C) Normalized gene expression plots of IFN-I genes in COVID-19 (top) and HIV-1 (bottom). (D) Network plot of key pathways correlated with IFN-I signaling in COVID-19 (left) and HIV-1^+^ patients (right). Size of each center corresponds to the number of genes present in the pathway. Genes are colored based on Log-2-fold change in expression in HIV-1^+^ versus COVID-19 patients. (E) Violin plot of the normalized gene expression of the actin polymerization pathway across each condition, split by selected cell subsets exhibiting high expression. (F) Violin plot of the normalized gene expression of the TLR signaling pathway across each condition, split by selected cell subsets exhibiting high expression.

IFN-I signaling plays pleiotropic roles during viral infection, including stimulating T cell survival, proliferation, and memory formation. We reasoned that the distinct or “modular” expression of IFN-I-associated genes during COVID-19 and HIV-1 could help explain some of the differences in these two viral infections. Using bicorrelation analysis on disease-specific IFN-I genes, we found a much higher number of top correlated genes and enriched pathways in COVID-19 compared to HIV-1 (Figure 6D). Notably, COVID-19 IFN-I correlated genes were enriched in MAPK signaling and interleukin signaling, which have been implicated in inflammation, thrombosis, and pulmonary injury and cytokine storm, respectively (Figure 6D, left). The ubiquitin-proteasome system (UPS) has been known to be a key target of viral manipulation (including SARS-CoV) in order to facilitate the production of viral proteins (Longhitano et al., 2020). As a result, proteasome inhibitors have been proposed for COVID-19 treatment (Longhitano *et al*., 2020). We also found an enrichment of proteasomal genes in COVID-19 patients, suggesting their unique role during SARS-CoV-2 infection (Lee et al., 2021).

Finally, we found an enrichment of the Rho GTPase metabolic pathway in COVID-19 patients (Figure 6D, left). Rho GTPases regulate a diversity of cellular processes, including cell migration and cell cycling, as well as modulating cytoskeletal rearrangement (Hodge and Ridley, 2016). The overlap between Rho GTPase and actin cytoskeleton genes confirmed this relationship. The actin cytoskeleton signature was especially pronounced in effector CD8^+^ T cells and antigen presenting cells in COVID-19 patients, indicating their preferential response (Figure 6E). GTPase activation has been found to contribute to immune cell activation and migration, as well as coagulation, often resulting in severe lung injury (Abedi et al., 2020a). In contrast, fewer significant pathways correlated with IFN-I signaling in HIV-1^+^ patients were all related to immune activation, including TLR signaling and CD28 co-stimulation (Figure 6F). Chronic viral antigen in HIV-1^+^ patients can induce chronic inflammation and constitutive TLR signaling, partly via LPS translocation which can lead to disease progression (Brenchley et al., 2006; Meier and Altfeld, 2007). Interestingly, we found that TLR signaling signature to be strongest in effector CD4^+^ and CD8^+^ T cells. Overall, our analysis demonstrates the differential IFN-I signaling following SARS-CoV-2 and HIV-1 infections and suggests that IFN-I signaling could play a more significant role in activating a greater diversity of immune cell functions in COVID-19 compared to HIV-1 (Lee and Shin, 2020; Schreiber, 2020).

### Metabolic differences between COVID-19 and HIV-1

The strong correlation of COVID-19 IFN-I signaling with Rho GTPase signaling suggested that enhanced IFN-I signaling in T cells could give rise to divergent metabolic profiles. Supporting this, two recent studies demonstrated that cellular metabolism is intimately linked to Rho GTPase activation and actin cytoskeleton organizations (Hu et al., 2016; Wu et al., 2021a). Our analysis also consistently revealed pathways associated with apoptosis and impaired metabolic function (Figure 5F). We hypothesized that SARS-CoV-2 and HIV-1 infections may also induce metabolically distinct signatures in immune cells. To investigate this, we built gene modules from the differentially enriched pathways we found and scored their expression. We found that immune cells in COVID-19 exhibited significantly lower mitophagy (Figure 7A, left), a process that maintains cellular homeostasis by removing damaged or dysfunctional mitochondria (Chen et al., 2020a) compared to immune cells in HIV-1 and healthy controls. Previous studies have reported that SARS-CoV-2 can activate the coagulation cascade in the blood, which could lead to a reduction in mitophagy (Ganji and Reddy, 2021). This altered rate of mitophagy forces cells to adopt apoptosis as an alternative, which could explain the elevated levels of apoptosis seen in COVID-19 cells compared to HIV-1 and healthy controls. However, we did not find a severity-dependent expression of mitophagy in COVID-19 patients (Figure 7A, right). We also confirmed the high expression of Rho GTPase activity in COVID-19 patients (Figure 7B, top left); while once again the expression was not stratified by disease severity (Figure 7B, top right), the trend was clear across every major cell type (Figure 7B, bottom). Finally, we found that while immune cells in COVID-19 patients upregulated the mTOR pathway compared to healthy controls, immune cells in HIV-1^+^ patients downregulated mTOR (Figure 7C, top). The mTOR pathway plays an important role in the response to viral infection by regulating cell proliferation and survival, as well as CD4^+^ T cell and B cell responses (Akbay et al., 2020; Ye et al., 2017). HIV-1 infection has been shown to interfere with mTOR signaling, usually resulting in diminished levels of mTOR expression in immune cells, particularly in CD4^+^ T cells (Akbay *et al*., 2020). We saw the same trend in CD4^+^ T cells of HIV-1^+^ patients (Figure 7C, bottom), which had the lowest mTOR expression across all cell types. These results suggest that elevated Rho GTPase and mTOR pathway activity and downregulated mitophagy is specific to COVID-19. Inhibitors to these pathways could be potential therapeutic targets for COVID-19 patients to alleviate the excessive activation commonly seen in severe cases.

**Figure 7:**
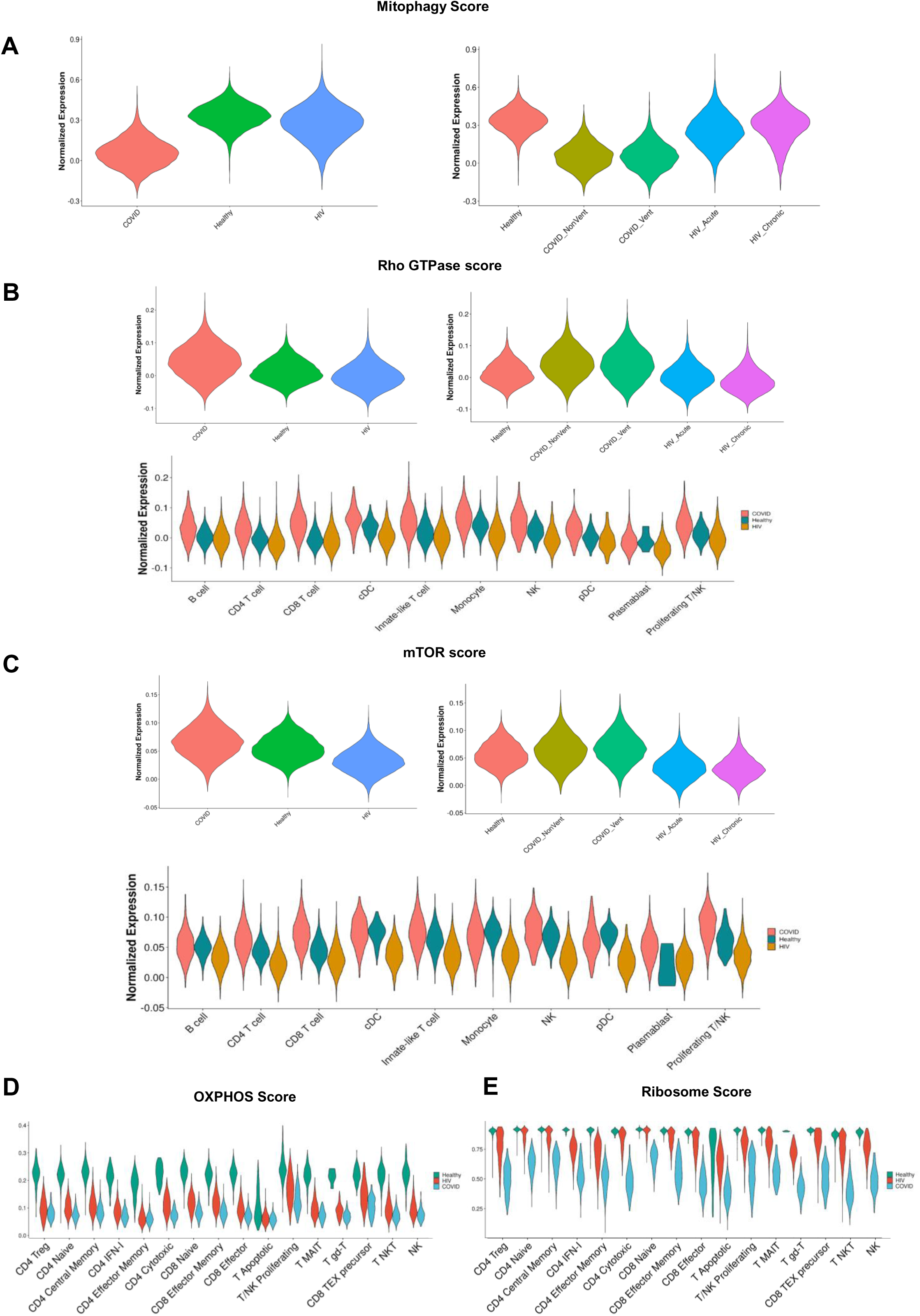
Metabolic differences associated with HIV-1 and COVID-19. (A) Violin plots of the normalized gene expression of the mitophagy pathway signature, split across diseases (left) and disease conditions (right). (B) Violin plots of the normalized gene expression of the Rho GTPase pathway signature, split across diseases (left), disease conditions (right), and major cell populations (bottom). (C) Violin plots of the normalized gene expression of the mTOR pathway signature, split across diseases (left), disease conditions (right), and major cell populations (bottom). (D) Violin plot of the normalized gene expression of the oxidative phosphorylation pathway across each condition, split by cell subset. (E) Violin plot of the normalized gene expression of the ribosomal pathway across each condition, split by cell subset.

We further examined metabolic signatures in T cells. We found specific downregulation of genes associated with autophagy, which is the process of degrading unnecessary cellular material. Since a subtype of autophagy is to selectively degrade damaged mitochondria, (i.e. mitophagy) our findings are consistent with previous reports that SARS-CoV-2 infection can cause the reduction of proteins responsible for autophagy initiation (Figure 5F). Additionally, general autophagy and specific mitophagy were both reported to be important for T-cell homeostasis, function, and differentiation. Deficiency in this process can lead to cell death, which may also explain the cell apoptosis and lymphopenia experienced by severe COVID-19 patients (Botbol et al., 2016; Gassen et al., 2021; Kovacs et al., 2012; Pua et al., 2009; Watanabe et al., 2014), and could be a potential therapeutic target. Interestingly, we also observed the shared downregulation of genes associated with ATP biosynthesis in both COVID-19 and HIV-1^+^ patients. Along with it, we found downregulation of genes associated with mitochondrial respiratory chain complex assembly in COVID-19 patients, and the comparable downregulation of OXPHOS in both groups (Figures 5F and 7D). Other studies have reported that mitochondria are affected during COVID-19 (Hoffmann et al., 2020; Singh et al., 2020). Prior studies have suggested that SARS-CoV-2 may hijack the host cell’s mitochondria, resulting in a reduction of ATP biosynthesis (Ganji and Reddy, 2021). Moreover, we found downregulation of genes associated with ribosome assembly in both groups, with a lower score in COVID-19 patients (Figure 7E). This result suggests that SARS-CoV-2 may also inhibit translation.

## Discussion

Since the COVID-19 pandemic, scRNA-seq has been extensively used to study the immune landscape of COVID-19 (Melms et al, 2021; Lee et al, 2020; Wen et al, 2020; Liao et al, 2020). While HIV-1 has been studied for almost 4 decades, the gene expression profile of HIV-1 infection at the single-cell level remains understudied. In particular, integration of scRNA-seq data across studies remains challenging and few attempts were made to integrate scRNA-seq data from COVID-19 and HIV-1^+^ patients. Here, we designed a consensus integration strategy that combined the advantages of deep-learning-based label transfer, molecular-profile-correlation-based label transfer and manual-supervised annotation methods that can be readily applied to scRNA-seq datasets. We leveraged the accuracy and portability of our method to generate a high-quality unified cellular atlas of the immune landscape of PBMCs from COVID-19 and HIV-1^+^ patients.

While manually supervised annotation offers domain-specific knowledge of known immune subset markers, a correlation-based method such as SingleR (Aran *et al*., 2019) complements it with unbiased comparisons with sorted a purified immune subset from healthy donors. However, both methods rely on the most typical and commonly known markers of different immune cell types, and neither incorporates disease-specific or context-specific knowledge from past studies of similar disease or context. This limitation can be well addressed with deep-learning based classification methods such as scANVI (Xu *et al*., 2021). The deep generative neural network can learn highly non-linear representations of each immune subset from the most finely-manually-annotated disease-specific scRNA-seq atlas and can leverage on the knowledge to classify new cells from the same disease in the same representation space. Nevertheless, overfitting can be a common issue for deep learning models, and expert knowledge is still required to keep the classification results in check. Thus, our strategy effectively consolidates the advantages of all three methods to annotate scRNA-seq data by 1) overcoming the subjectivity of manual annotation, 2) leveraging existing knowledge of cell phenotypes to quickly and accurately assign labels with a trained model, and 3) validating classified labels with biologically relevant markers.

Besides integration of COVID-19 and HIV-1 PBMC data, we anticipate our integration strategy can be easily adapted for integration of scRNA-seq data from different tissues, organs, or diseases. While we highlighted the integration of manual, correlation-based, and deep-learning-based annotation methods, there is flexibility for the specific software used in each method. The software we used in this study, namely Seurat, SingleR and scANVI, are all publicly available and highly rated across multiple benchmarking studies (Abdelaal *et al*., 2019; Huang et al., 2021; Krzak et al., 2019), so the current implementation can be adapted as is. One potential limitation is that there may not be high quality reference data for training the deep learning model for certain context or disease. However, we envision it will be increasingly easy to overcome this, given multiple ongoing efforts to make large atlases of specific tissues, organs, and diseases easily accessible such as Azimuth (Hao et al., 2021) and the human protein atlas (Uhlen et al., 2017). Using our integration strategy, we identified 27 different cell types, consisting of 5 B cell subsets, 2 DC subsets, 4 monocyte subsets, 7 CD4^+^ T cell subsets, 8 CD8^+^ T cell subsets, and 1 NK cell subset. We also conducted detailed comparison within each immune compartment between each disease against healthy control as well as between the two diseases to identify the key common and differential regulatory pathways.

We found a consistent inflammatory signature highlighted by IFN-I and cytokine-mediated signaling among innate immune cells in both diseases (Bieberich et al., 2021; Hasan et al., 2021; Liu *et al*., 2021). However, we also discovered that the types and frequencies of cellular communications among immune cells can be very different between COVID-19 and HIV-1 patients. This allowed us to identify the disease-specific inflammatory and cytotoxic molecules that drive the innate immune response in either disease. Interestingly, we found an enrichment of inhibitory interactions mediated by *CTLA4* and *HAVCR2* that were unique to COVID-19 patients, which could be out of necessity to curb the heightened inflammation present in severe COVID-19. Further experiments are necessary to validate these hypotheses.

Consistent with prior literature, we found a strong humoral immune response in both COVID-19 and HIV-1^+^ patients (Baum, 2010; Wu et al., 2021b) driven by plasmablast maturation and activation. While we could not evaluate the functionality of the antibody repertoire by single-cell RNA-seq analysis, we were able to demonstrate that the HIV-1 repertoire was much less diverse compared to the COVID-19 repertoire. Our study is aligned with the vaccine efficacy results against both viruses. Multiple SARS-CoV-2 vaccines have shown efficacy due in part to their ability to generate broadly protective neutralizing antibodies, but this is not the case for candidate HIV-1 vaccines (Baden et al., 2021; Baum, 2010; Dangi et al., 2021a; Dangi *et al*., 2021b; Goel et al., 2021a; Mercado et al., 2020; Polack et al., 2020; Sanchez et al., 2021; Turner et al., 2021). Repertoire mapping also allowed us to pinpoint high-frequency as well as overlapping combinations and could inspire antibody-based therapeutics to treat comorbid patients. Additionally, we found the COVID-19-specific IGKV1-39/IGHV2-26 combination to have significant enrichment among our patients, which could pose as a promising therapeutic candidate.

IFN-I can play a double-edged role during viral infection; while it restricts viral replication and viral antigen expression in addition to modulating the antigen-specific CD8^+^ T cell response (Moseman et al., 2016; Palacio et al., 2020), overactive IFN-I signaling can contribute to immune dysfunction, non-canonical inflammasome activation, and pyroptosis (Kopitar-Jerala, 2017; Teijaro et al., 2013; Wilson et al., 2013). Previous studies comparing the immune response in COVID-19 and influenza have investigated the role of IFN-I signaling in both driving the antiviral response and disease progression (Galani et al., 2021; Lee *et al*., 2020; Nguyen et al., 2021). While a positive effect of IFN-I has been defined in the immune response to influenza, the role of IFN-I during severe COVID-19 remains unclear. In acute HIV-1 infection, IFN-I signaling has been generally characterized as beneficial (Abraham et al., 2016; Lavender et al., 2016; Sandler et al., 2014; Wang et al., 2017). However, during the late stages of chronic viral infection, IFN-I signaling shifts toward a pathogenic role by contributing to systemic inflammation (Soper et al., 2017; Teijaro *et al*., 2013; Utay and Douek, 2016; Wilson *et al*., 2013). However, the precise role of IFN-I at the single-cell level in COVID-19 (which results in an acutely controlled infection) and HIV-1 (which results in a chronic infection) is still unclear.

We identified IFN-I signaling to be a key pathway induced across various immune cells. IFN-I is critical to prime innate and adaptive immune responses during both SARS-CoV-2 and HIV-1 infection, as well as limiting viral replication and promoting effector cell function (Schreiber, 2020; Sugawara et al., 2019). As a result, IFN-I therapy has been proposed for both diseases. While IFN-I signaling was upregulated in both COVID-19 and HIV-1 patients relative to healthy controls, our analysis suggests a more robust role of IFN-I in COVID-19. We found that IFN-I signaling in COVID-19 is more intimately tied to important cellular functions such as cell signaling, motility, and cytokine secretion. In support of our findings, previous studies have found that exposure to IFN-I results in upregulation of MAPK signaling cascades (Zhao et al., 2011). While MAPK signaling regulates important functions such as cellular proliferation and survival, further studies are needed to investigate whether IFN-I mediated MAPK signaling in COVID-19 contributes to antiviral immune response or apoptosis (Zhang and Liu, 2002). Dysregulation of actin cytoskeleton following viral infection activates RLR signaling and downstream IFN-I signaling, which could explain the origin of the IFN-I response in COVID-19 patients (Trono et al., 2021). Previous studies have reported an antagonistic relationship between IFN-I and IL-1, the prototypical proinflammatory cytokine (Guarda et al., 2011; Mayer-Barber and Yan, 2017). Interestingly, we found that IFN-I signaling in COVID-19 patients is highly correlated with immune-activating cytokine signaling pathways such as IL-2, IL-16, and IL-17, which could provide novel insights on the coregulatory relationship of IFN-I with other effector cytokines. In contrast, we found a much narrower scope of highly correlated genes and pathways in HIV-1^+^ patients, including the CD161^+^ CD8^+^ T cell signature and TLR signaling. CD161^+^ CD8^+^ T cells represent a subset of innate-like memory CD8^+^ T cells that feature elevated levels of cytotoxicity, cytokine production, and survival, in addition to providing antigen-specific protection against viruses such as HBV, CMV, and influenza (Fergusson et al., 2016; Konduri et al., 2020). While CD161^+^ CD8^+^ T cells have not been well characterized in the context of HIV-1, CD161^+^ CD8^+^ T cells have demonstrated increased sensitivity to IFN-I stimulation, synergizing with TCR/CD3 activation to trigger high cytotoxicity and cytokine production (Pavlovic et al., 2020). Thus, the IFN-I driven CD161^+^ CD8^+^ T cell response can be an important antiviral mediator during HIV-1 infection. Additionally, the correlation of TLR signaling with IFN-I signaling is expected, since TLR triggering induces downstream IFN-I responses and subsequent induction of interferon-stimulated genes (Uematsu and Akira, 2007; Wang et al., 2019). However, the specific enrichment of TLR signaling in HIV-1^+^ patients could suggest a disease-specific driver of immune activation. Stimulation of TLRs can result in latency reversal (Macedo et al., 2019; Meås et al., 2020), which in turn activates IFN-I production, thereby contributing to chronic inflammation. Overall, our findings show that while IFN-I response is robust in both diseases, they are tied to drastically different biological functions in HIV-1 compared to COVID-19, with the latter featuring a much more diverse spectrum of cellular responses. These insights are important to consider given the recent proposals to utilize IFN-I as a potential treatment for COVID-19. In agreement with our findings, recent analyses integrating genome-wide association study (GWAS) and transcriptome-wide association study (TWAS) suggested an important role of IFN responses in determining the COVID-19 severity (Pairo-Castineira *et al*., 2021) .

We reason that our gene expression comparison could elucidate pathways that could be targeted for the treatment of COVID-19 or HIV-1. In particular, our analyses corroborated the contribution of various molecular pathways that regulate COVID-19 pathophysiology, many of which are already considered for COVID-19 treatments. For instance, we found JAK-STAT signaling, IL-4 signaling, and IL-6 signaling to be enriched in COVID-19 patients, and interestingly, all of these pathways have been targeted for the treatment of COVID-19. Three JAK inhibitors, namely Baricitinib, Tofacitinib, and Ruxolitinib have been used to treat COVID-19 patients by reducing excessive inflammation, among which Baricitinib and Tofacitinib are recommended for hospitalized patients who require high-flow oxygen or noninvasive ventilation according to NIH COVID-19 Treatment Guidelines (Satarker *et al*., 2021). In addition, Dupilumab, an IL-4Rα inhibitor, was also reported to be useful for treating COVID-19 patients (Thangaraju et al., 2020). Furthermore, IL-6R inhibitors Sarilumab and Tocilizumab were also shown to be beneficial for COVID-19 patients and were recommended for use in hospitalized patients who require supplemental oxygen, high-flow oxygen, noninvasive ventilation, or invasive mechanical ventilation by NIH COVID-19 Treatment Guidelines. Interestingly, we also found COVID-19 specific enrichment of *MAP3K1*, which activates the JNK and ERK pathway. Since JNK inhibition were reported to preclude the development of persistent SARS-CoV infections (Mizutani et al., 2005) and MEK1/2 inhibition can diminish the production of viral progeny of coronavirus (Cai et al., 2007), inhibitors targeting both JNK and ERK pathway have the potential to be used for COVID-19 treatment. Besides, our findings suggest that COVID-19-specific IFN-I correlated genes were enriched in MAPK signaling, and p38 MAPK inhibition was previously reported to reduce human coronavirus HCoV-229E viral replication in human lung epithelial cells (Kono et al., 2008). Therefore, inhibitors targeting MAPK signaling also have the potential to be used for treating COVID-19 patients. We also found the IFN-γ/IFN-γ receptor interaction between CD8^+^ T cells and APCs to be enriched in HIV-1^+^ patients. While IFN-γ production in the acute phase of HIV-1 infection can help curtail infection, it can also contribute to persistent inflammation and tissue damage during the chronic disease (Roff et al., 2014). We found this interaction to be specifically active between CD8^+^ T cells and dendritic cells and monocytes in both acute and chronic HIV-1^+^ patients. Thus, molecules aimed to stimulate or inhibit IFN-γ signaling in specific cell types could help address HIV-1 pathogenesis at different stages of disease. In addition, our studies independently highlighted the significance of JAK and IFN-I in SARS-CoV-2 infection which was suggested by recent human GWAS and TWAS results (Pairo-Castineira *et al*., 2021).

Notably, our analysis also revealed disease-specific altered metabolism profiles. We characterize a decrease in OXPHOS and ribosome biogenesis in response to both SARS-CoV-2 and HIV-1 infection. Virus-induced reduction of OXPHOS has been previously characterized in other diseases and could be a result of oxidative stress triggered by mitochondrial clustering (Khan et al., 2015). Viral hijacking of ribosomal function is also crucial to viral replication and survival in the host (Li, 2019). Disruption of these viral interactions could be advantageous for COVID-19 and HIV-1 treatment. We also unveiled molecular metabolic pathways which could also be targeted for therapy. For example, we found enriched proteasomal genes in COVID-19 patients. It has been proposed that proteasome inhibitors may be a possible therapy for COVID-19, since proteasome inhibitor may interfere with the viral replication processes and reduce the cytokine storm associated with various inflammatory conditions (Longhitano *et al*., 2020). Consistent with our observation of the upregulation of Rho GTPase in COVID-19 patients; the plausibility of using Rho kinase inhibitors to treat COVID-19 has been discussed, as they can restore the activity and level of ACE2 which is inhibited by SARS-CoV-2 without increasing the risk of infection (Abedi et al., 2020b). Although remained to be investigated in immune cells, a recent study demonstrated that small GTPase RhoA activation drives increased cellular glycolytic capacity (Wu *et al*., 2021a) which is typically associated with reduced mitochondrial metabolism, in agreement with the upregulation of Rho GTPase and disrupted mitochondrial function in COVID-19 patients. Moreover, Rho GTPases have been linked to additional key metabolic controls such as mTOR signaling pathways. (Mutvei et al., 2020; Senoo et al., 2019). Finally, we found COVID-19 specific upregulation of the mTOR pathway, and thus its inhibitors may also be used for treatment, since mTOR inhibitors can adjust T cells by induction of autophagy without apoptosis, reduce viral replication, restore T-cell function, and decrease cytokine storm (Mashayekhi-Sardoo and Hosseinjani, 2021).

In conclusion, our study provides a comprehensive comparison of the immunological landscape of SARS-CoV-2 and HIV-1 infections in humans. The high resolution of single-cell RNA sequencing, diversity of patient samples, and large dataset allowed us to unveil important shared and disease-specific features that offer insight into the next generation of antiviral treatments. Through cell type-specific analysis, we found a common enrichment of activated B cells and plasmablasts, inflammatory monocyte and effector T cell subsets, and cytokine signaling that appear to drive the antiviral response to SARS-CoV-2 and HIV-1. We also found dendritic cells and monocytes to be highly interactive with adaptive immune cells in both diseases, but found that innate cells in COVID-19 appear to be more capable of immunosuppressive function through CTLA-4 and TIM-3-mediated interactions. We also found that the cytokine response was more diverse in COVID-19 patients, which is highlighted by IL-2, IL-4, and IL-20 signaling, while HIV-1^+^ individuals primarily exhibited high levels of NF-kB signaling. Our analysis corroborated pathways in COVID-19 patients that have already shown therapeutic benefits, but further *in vitro* and *in vivo* experiments are necessary to measure the contribution of other molecular pathways (such as Rho GTPase and IL-2 signaling) that also appear to be distinctively enriched. Overall, our study provides a roadmap to help develop novel drugs to treat COVID-19 and HIV-1 infections.

## Methods

### Preprocessing, integration, and clustering

Raw single-cell count matrices were collected from publicly available sources (PBMCs from a Healthy Donor: Whole Transcriptome Analysis, 2020; Kazer *et al*., 2020; Wang *et al*., 2020; Wilk *et al*., 2020) and merged. We performed quality control and downstream analysis using the Seurat package (v4.0.4) (Stuart et al., 2019). We removed cells with greater than 15,000 UMIs or fewer than 500 UMIs, as well as greater than 20% mitochondrial reads per cell, resulting in a total of 115,272 cells. We performed log-based normalization with the “NormalizeData” function with the “LogNormalize” parameter and selected the top 10,000 variable features with the “vst” parameter using “FindVariableFeatures”. We scaled and centered the count matrix using the “ScaleData” function and supplied “percent.mito” as a latent variable to regress out the effect of percentage mitochondrial reads. We performed principal component analysis (PCA) on the top 100 PCs using the “RunPCA” function. To remove study-specific batch effects, we performed integration across each patient using the Harmony algorithm (v0.1.0) (Korsunsky et al., 2019) on the top 50 PCs with the “RunHarmony” function. We then performed Uniform Manifold Approximation and Projection (UMAP) reduction using the “RunUMAP” function on the top 50 PCs with “min.dist” = 0.1 and “n.neighbors” = 20. We ran the “FindNeighbors” function on the top 50 Harmony dimensions, then performed Louvain clustering using the “FindClusters” function with a resolution of 0.3. We annotated the clusters using known cell type-specific markers, resulting in a total of 19 cell types, including 7 CD8^+^ T cell subtypes, 3 Monocyte subtypes, and 4 CD4^+^ T cell subtypes, and added the labels to the main object.

### Cell subset annotation

For manual annotation, we subsetted the three major cell populations (T cells + NK cells, B cells + Plasmablasts, and Dendritic cells + Monocytes) and separately performed normalization, scaling, feature selection, PCA, integration, UMAP, and clustering. For reference-based annotation, we utilized the “SingleR” method from the SingleR package (v1.4.1) (Aran *et al*., 2019) using data from Monaco et.al (Monaco et al., 2019) and default parameters, and transferred the fine and coarse labels to the main object. In total, we found 7 major cell types and 27 subtypes with SingleR. For deep learning-based annotation, we used the scANVI package (v0.7.0) from the scvi-tools library (Xu *et al*., 2021) to train a deep generative model using reference data from Ren et. al. (Ren et al., 2021). We first merged the raw counts from our object data with raw counts from the reference into a combined AnnData object. We normalized and logarithmized the matrix with the Scanpy package (Wolf et al., 2018) (v1.4.5) using the “normalize_total” method with “target_sum” = 10,000 and “log1p” method. We found highly variable genes using the “highly_variable_genes” method with “flavor” = “seurat_v3” and “n_top_genes” = 4000. In order to improve the accuracy of the model, we performed hierarchical clustering on the reference data and merged labels that fell under a common hierarchy, resulting in 32 total labels: 5 B cell subsets, 3 DC subsets, 4 Monocyte subsets, 7 CD4^+^ T cell subsets, and 8 CD8^+^ T cell subsets (Figures S1A and S1B). We supplied the resulting labels to the reference data. We subsampled approximately 500 cells from each cell subset from the reference data to act as the training set and built the model with a latent dimensionality of 30 and 2 hidden layers using the “model.SCANVI” method. We then trained the model using 300 passes for semi-supervised training using the “train” method. We obtained the labels using the “predict” method and transferred the labels to the main object. Our resulting model had an overall accuracy of 76% and a median F1 score of .78, which was significantly higher than the reported accuracy of existing methods such as LAmDA, scmapcluster, and LDA (Abdelaal *et al*., 2019) (Figures S1C and S1E).

### Consensus annotation

Generation of consensus markers was performed using the following steps:

1. Compare manual and SingleR labels. If they are identical, leave the label as-is.
2. If one label is at higher resolution (i.e. is a subset of the other), assign the higher resolution label.
3. If the two labels are inconsistent, subset out and pool with similarly inconsistent labels. Plot gene expression using markers of either label type. Assign the label with corresponding marker expression (Consensus 1).
4. Repeat 1-3 using Consensus 1 and scANVI labels.

### Cell type composition comparison

We computed frequencies of each cell type for each patient and performed Wilcoxon rank-sum tests on the medians to find significantly different compositions between pairs of patient types (HIV, COVID-19, and healthy).

### Cluster purity assessment

We utilized the ROGUE package (Liu *et al*., 2020) (v1.0) to assess purity of clusters determined by consensus labels. We calculated the expression entropy of each gene using the “SE_fun” method with “span” = 1.0. We calculated the ROGUE value of each consensus label across each patient using the “CalculateRogue” function with “platform” = “UMI”.

### Differential gene expression analysis and gene set enrichment analysis (GSEA)

To compare the relative similarities and differences of HIV-1 and COVID-19 gene expression, we performed differential gene expression analysis for either disease with respect to healthy controls. Differentially expressed genes were determined using a Wilcoxon Rank Sum test with Seurat’s “FindMarkers” function with the parameters “logfc.threshold” = 0 and “min.pct” = .1. P values were adjusted based on bonferroni correction. We denoted differentially expressed genes (DGEs) with average log-2-fold change greater than 1 or less than -1 as differentially upregulated or downregulated, respectively. We performed GSEA on DGEs using the clusterProfiler package (v3.18.1) (Yu et al., 2012) with the “GSEA” function using default parameters using pathways from the MSigDB database (Subramanian et al., 2005).

### B cell chain analysis

We determined the frequency of heavy chain/light chain combinations using a method adopted from Melms (Melms *et al*., 2021). We filtered B cells and Plasmablasts to only the cells that expressed both heavy chain (IGVH) and light chain (IGVL) genes. These consisted of genes beginning with IGHG, IGHM, IGHA, or IGHE for heavy chain genes, and IGLV or IGKV for light chain genes. We then counted the number of mRNA transcripts for each IGVH and each IGVL gene expressed on a per-cell basis, then assigned the most highly expressed IGVH and IGVL genes to be that cell’s IGVH-IGVL pairing. We aggregated the number of cells that expressed each IGVH-IGVL pairing and plotted the frequency of each combination in a heatmap (Figures S4A-D). If that pairing only appeared in HIV-1^+^ or COVID-19 patients, we assigned it to its respective disease. Otherwise, we classified as “shared”.

### Receptor-ligand analysis

To infer the putative receptor-ligand interactions between pairs of cell types, we utilized CellPhoneDB (Efremova *et al*., 2020). We first normalized raw count matrices to counts per 10,000 for each patient. We then performed CellphoneDB separately for each patient using the statistical method and default parameters, while supplying labels as the metadata for the 10 broad cell types. This was done to maintain biological accuracy, as feasible ligand-receptor interactions are only meaningful when measured within a given patient. We filtered out all ligand-receptor pairs with negative values, then merged interactions from patients of the same disease, treating each ligand-receptor/cell type combination as a unique interaction while preserving directionality (i.e. Monocyte-NK is unique from NK-Monocyte). This was done to capture the full spectrum of possible interactions across cell types. We averaged expression values and p-values for each interaction across patients. We repeated this process for all 27 consensus labels.

### IFN-I correlation analysis

We first compiled genes belonging to MSigDB pathways including the term “Type-I Interferon Signaling) into an IFN-I gene list. We then found differentially expressed genes between HIV-1^+^ and COVID-19 patients using the previously mentioned parameters and filtered them to keep only IFN-I related genes. We scored each gene module using the Seurat function “AddModuleScore”. To perform correlation analysis, we first used the SuperCell package (Okhotnikov et al., 2016) to group cells from each batch into supercells of 100 cells each using 5 K-Nearest-Neighbors and 2000 variable genes and combined the resulting gene expression matrices from common diseases together. We then ran the “bicor” function from the WGCNA package (v1.70) (Langfelder and Horvath, 2008) on each gene belonging to the COVID-19 IFN-I module with each gene present in the COVID-19 supercell matrix, and extracted the top correlated genes (>.65). This was repeated for the HIV-1 IFN-I module and supercell matrix. We then performed GSEA enrichment on each set of top correlated genes.

## Acknowledgements

This work was supported by NIH New Innovator award <u>1DP2AI144245</u> (J.H.), NSF Career award <u>1653782</u> (J.H.), and the Third Coast Center for AIDS Research (CFAR), an NIH funded center (P30 AI117943). T.P. is supported by NHLBI T32HL007381.

## Author Contributions

T.P and G.C. designed the original concept for the project. T.P., G.C., and Y.Z. performed analyses. T.P., G.C., and E.T. performed interpretation of results. J.H. conceived and supervised the project. T.P., G.C., and E.T. wrote the manuscript while J.H. edited the manuscript with inputs from P.P.M and Y.F.

## Supplemental Figure Legends

**Figure S1:**
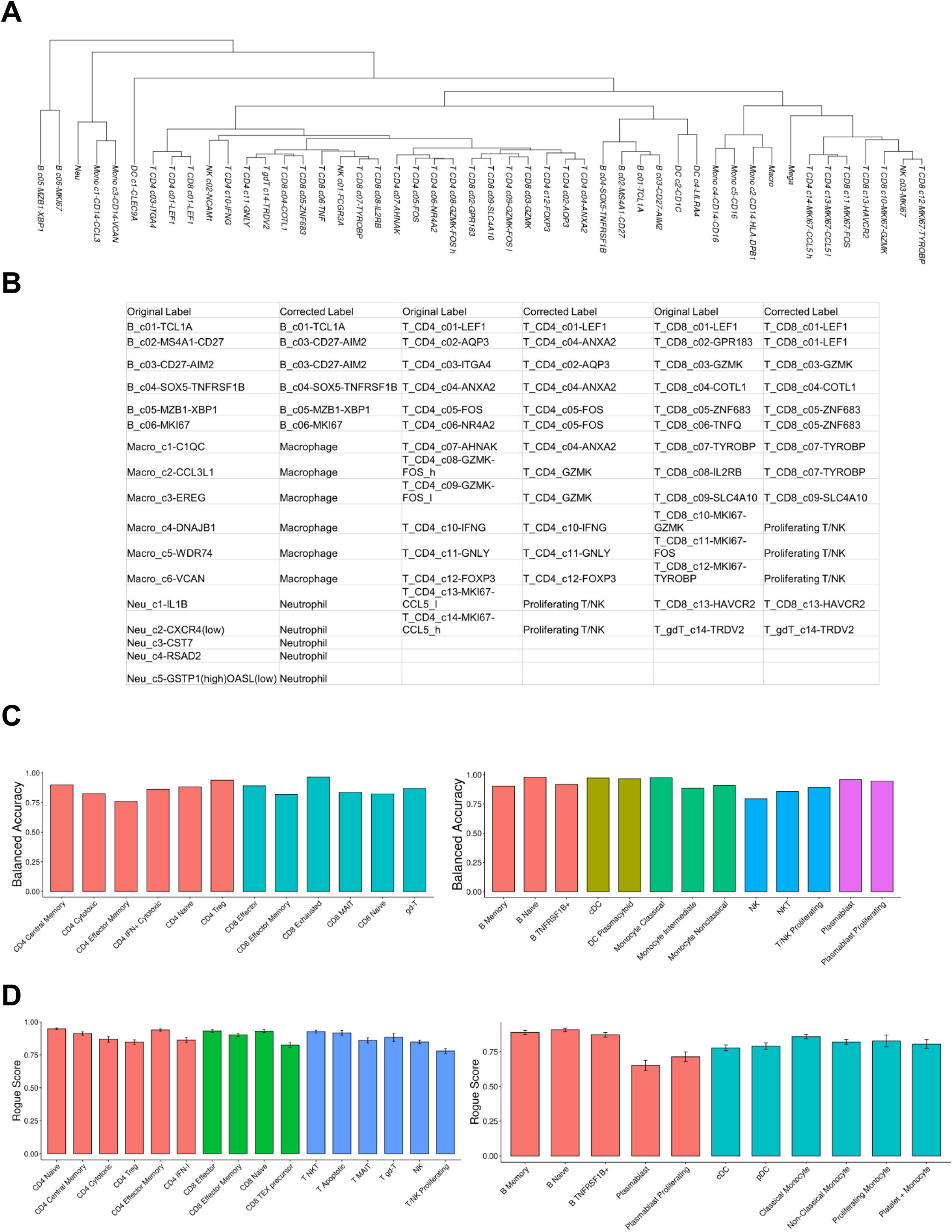
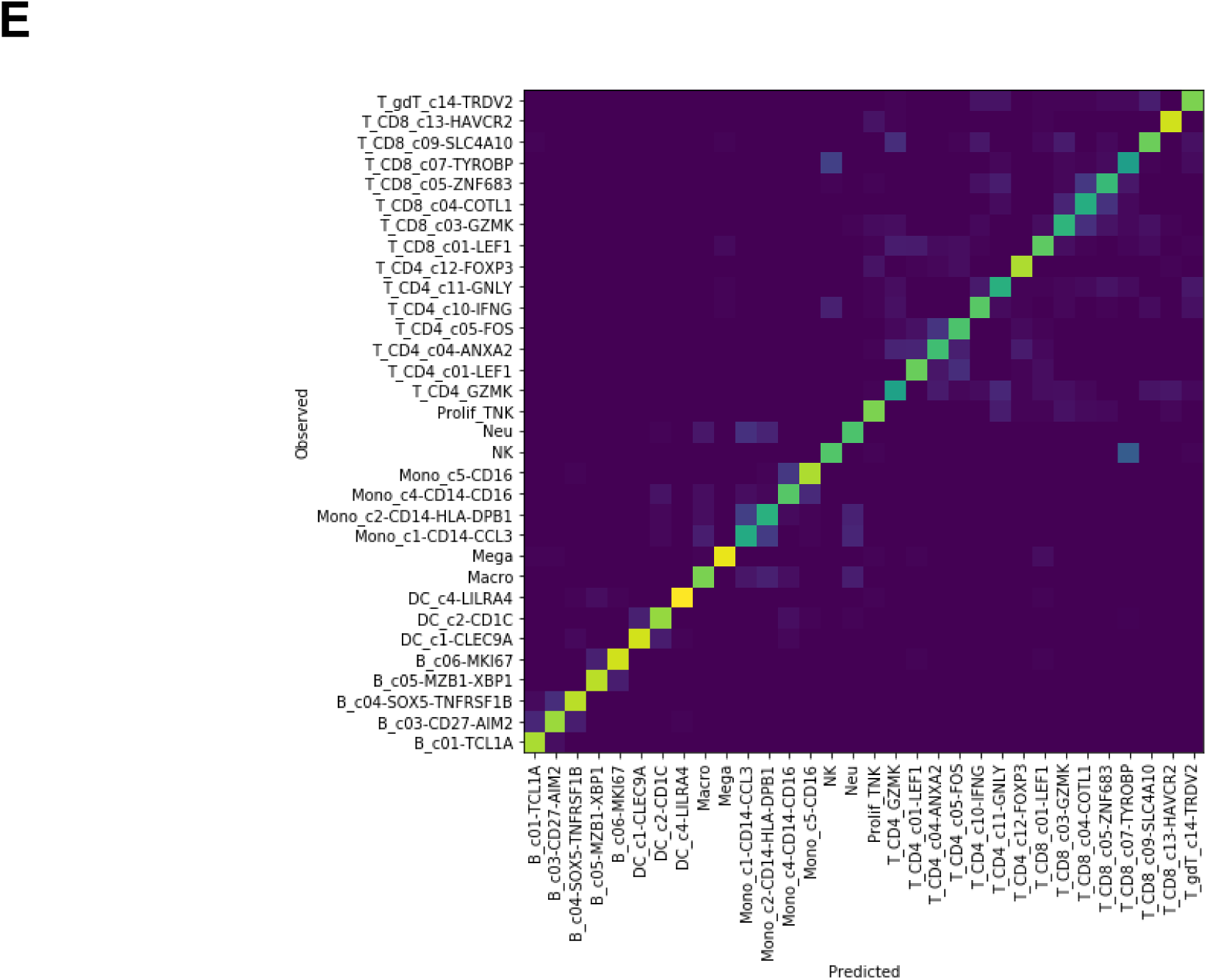
Assessment of scANVI training and consensus clustering. (A) Hierarchical clustering of original labels derived from Ren Cell 2021. (B) Table depicting the merging of labels from Ren Cell 2021 that were used for scANVI training. (C) Balanced accuracies of scANVI-derived labels compared against their original label from Ren Cell 2021. (D) ROGUE scores of final consensus labels. Error bars denote variation across patients. (E) Confusion matrix comparing the observed labels provided by scANVI (Y axis) and predicted labels (X axis)

**Figure S2:**
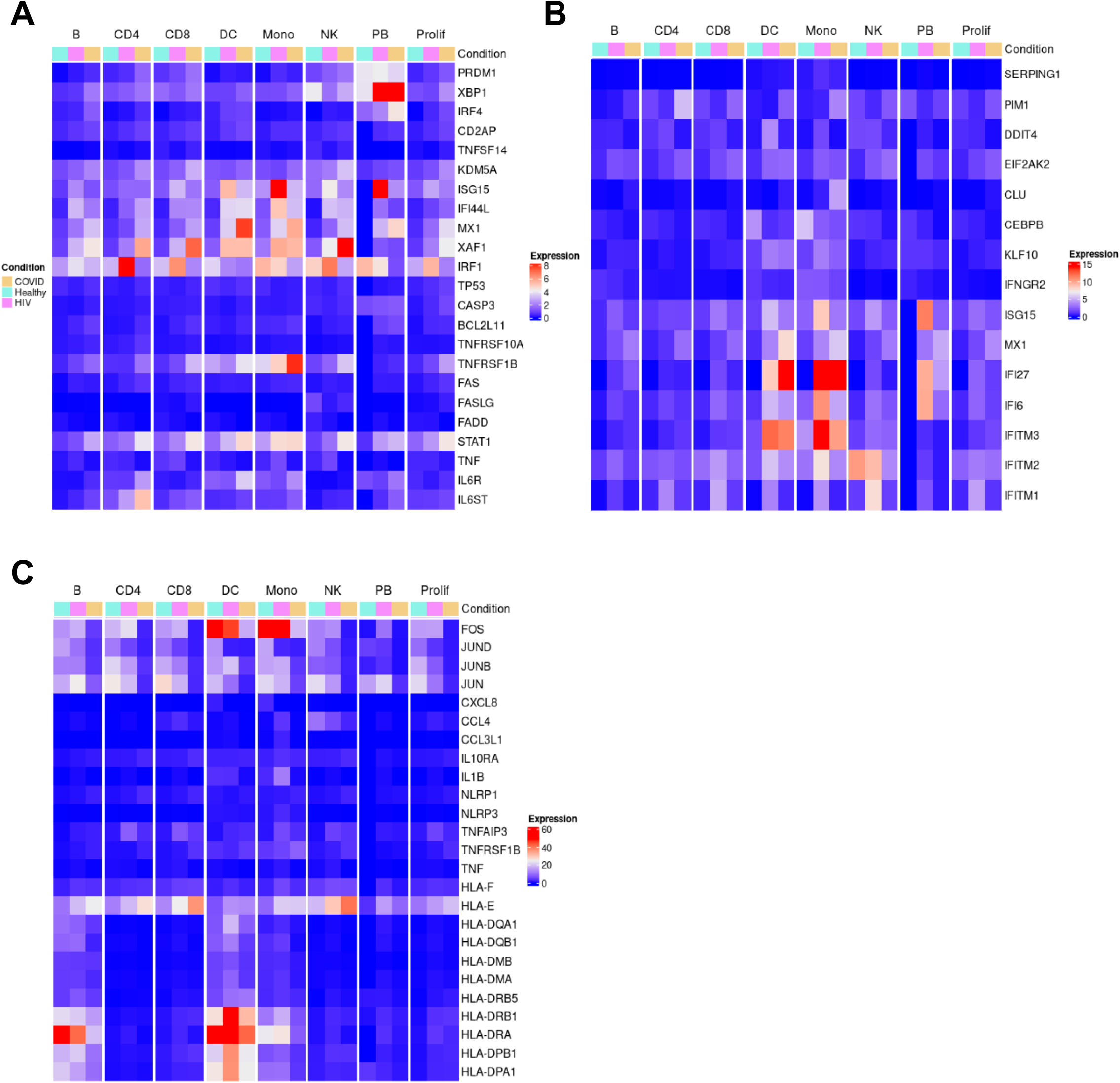
Gene expression heatmaps across COVID-19, HIV-1^+^, and healthy patients. (A) Heatmap of selected genes found to be upregulated in COVID-19 patients from Zhu Immunity 2020 (Zhu *et al*., 2020) plotted across cell types and patient conditions. (B) Heatmaps of selected genes found to be upreguated in COVID-19 patients from Xu Cell Discovery 2020 (Xu et al., 2020) plotted across cell types and patient conditions. (C) Heatmaps of selected genes found to be downregulated in COVID-19 patients from Xu Cell Discovery 2020 (Xu *et al*., 2020) plotted across cell types and patient conditions.

**Figure S3:**
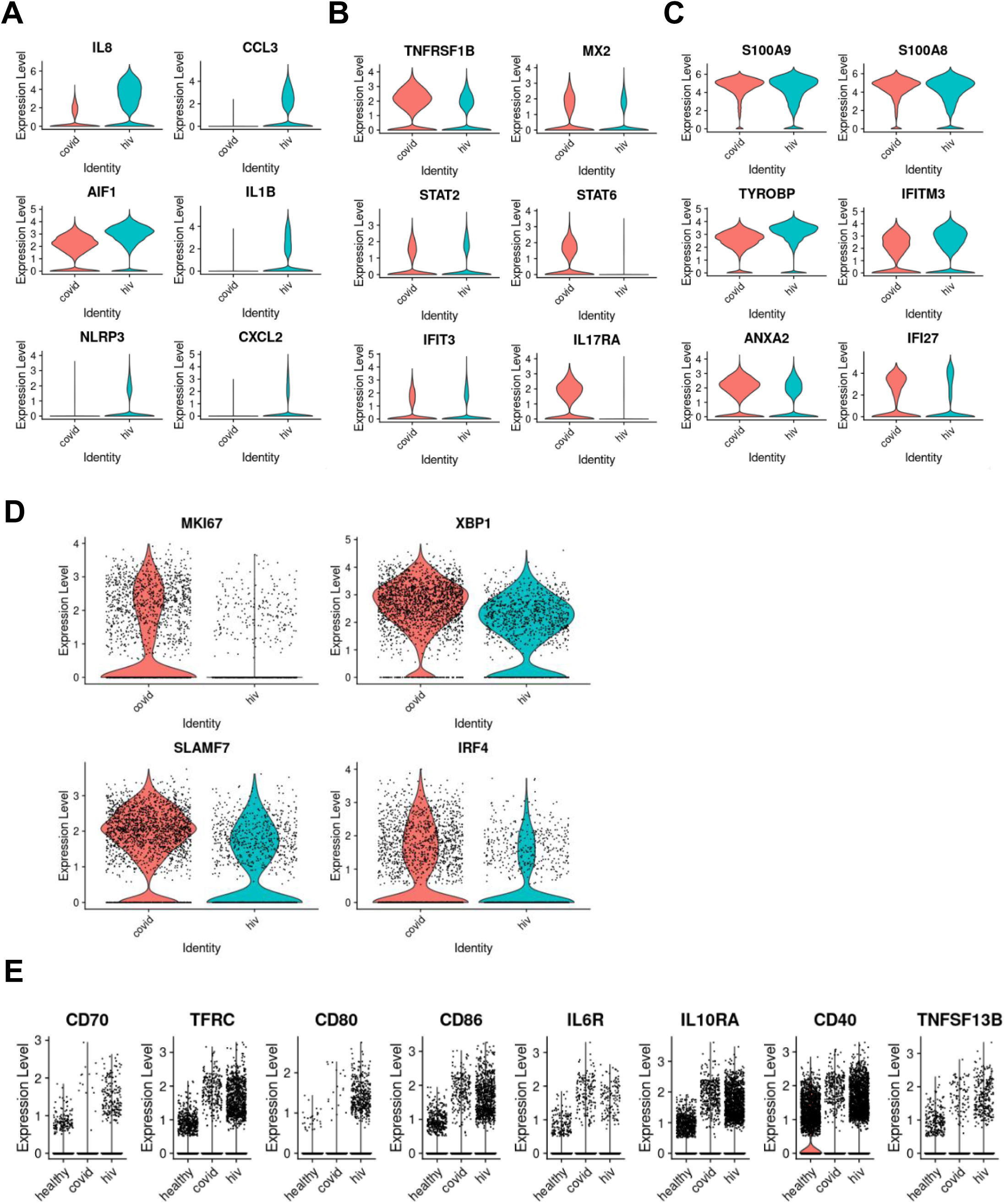
Violin plots of gene expression across COVID-19, HIV-1^+^, and healthy patients. (A) Violin plots of selected genes upregulated in monocytes from HIV-1^+^ patients compared to monocytes from COVID-19 patients. (B) Violin plots of selected genes upregulated in monocytes from COVID-19 patients compared to monocytes from HIV-1^+^ patients. (C) Violin plots of selected genes jointly upregulated in monocytes from COVID-19 and HIV-1^+^ patients. (D) Violin plots of selected genes in plasmablasts from COVID-19 and HIV-1^+^ patients. (E) Violin plots of selected genes in B cells from COVID-19 and HIV-1^+^ patients.

**Figure S4:**
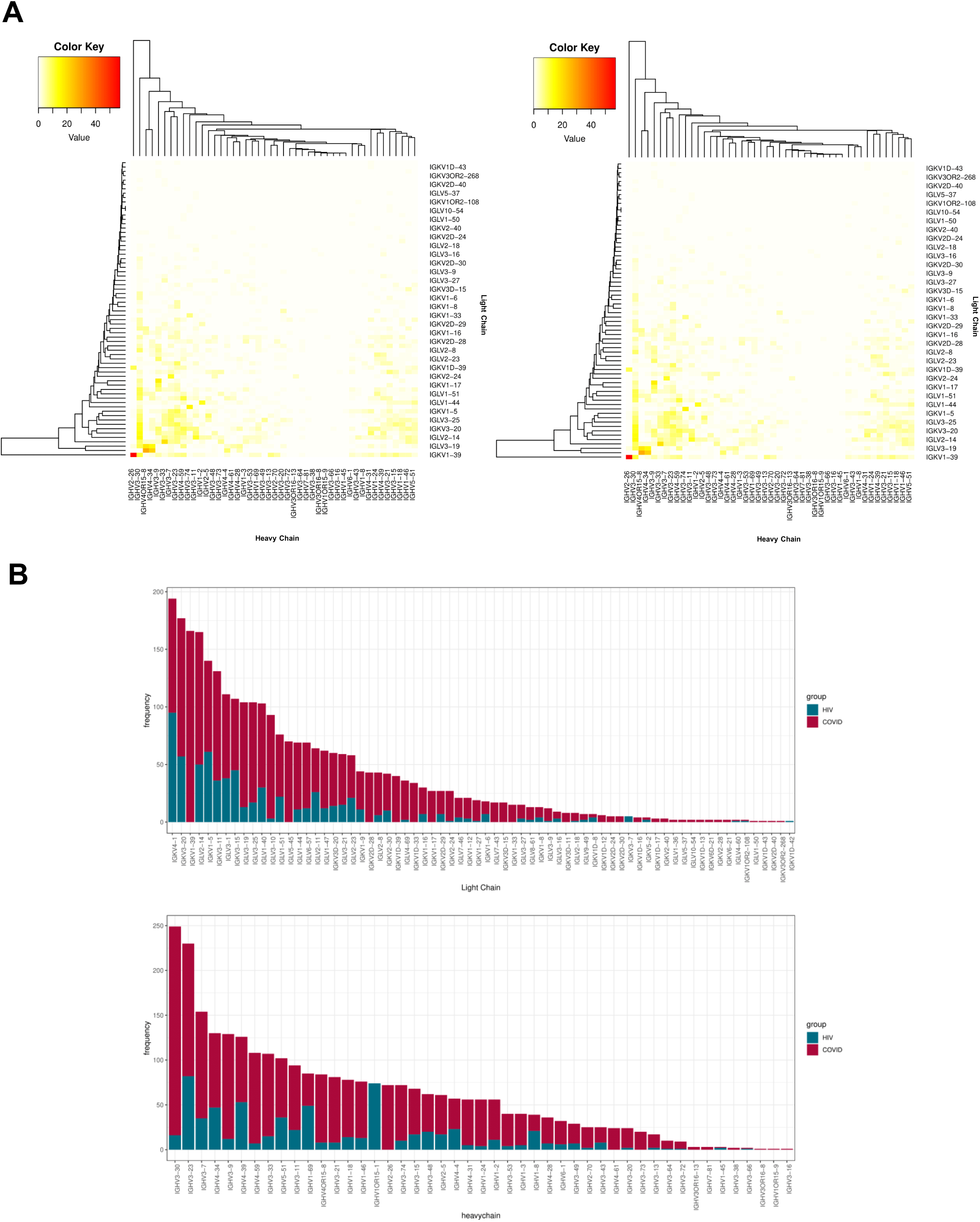
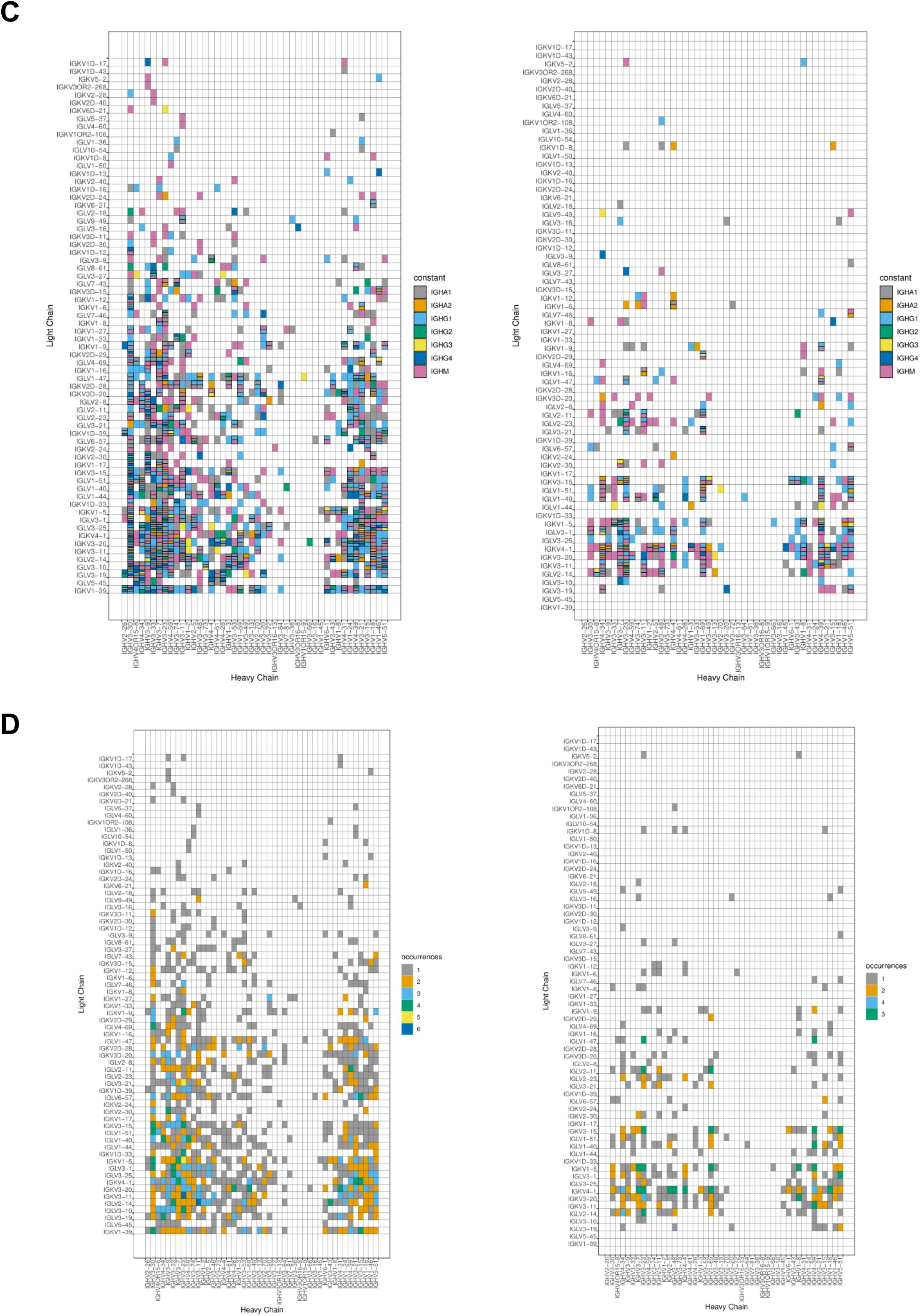
Immunoglobulin combinations in B cells and Plasmablasts. (A) Heatmap of the frequencies of variable heavy (X axis) and light (Y axis) chains in B cells and plasmablasts in COVID-19 (left) and HIV-1^+^ (right) patients. Color indicates number of cells expressing the specific heavy/light chain pairing. Hierarchical clustering was performed on average linkage. (B) Barplots of the frequency of each light chain (top) and heavy chain (bottom) compared across COVID-19 and HIV-1^+^ patients. (C) Heatmap of heavy chain (X axis) and light chain (Y axis) combinations colored by constant region in COVID-19 (left) and HIV-1^+^ (right) patients. (D) Heatmap of all heavy chain (X axis) and light chain (Y axis) combinations colored by frequency expressed in patients in COVID-19 (left) and HIV-1^+^ (right) patients.

**Figure S5:**
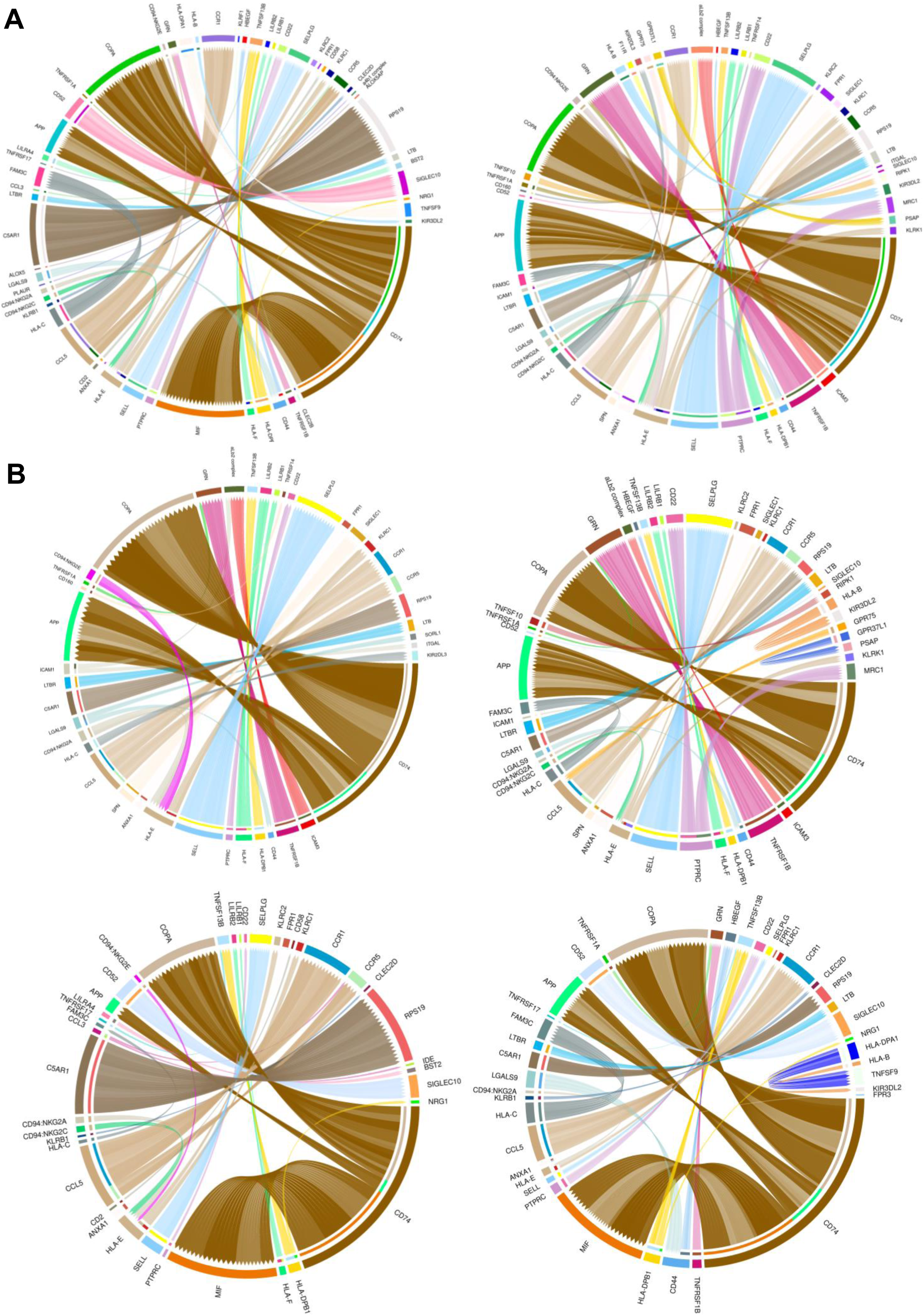
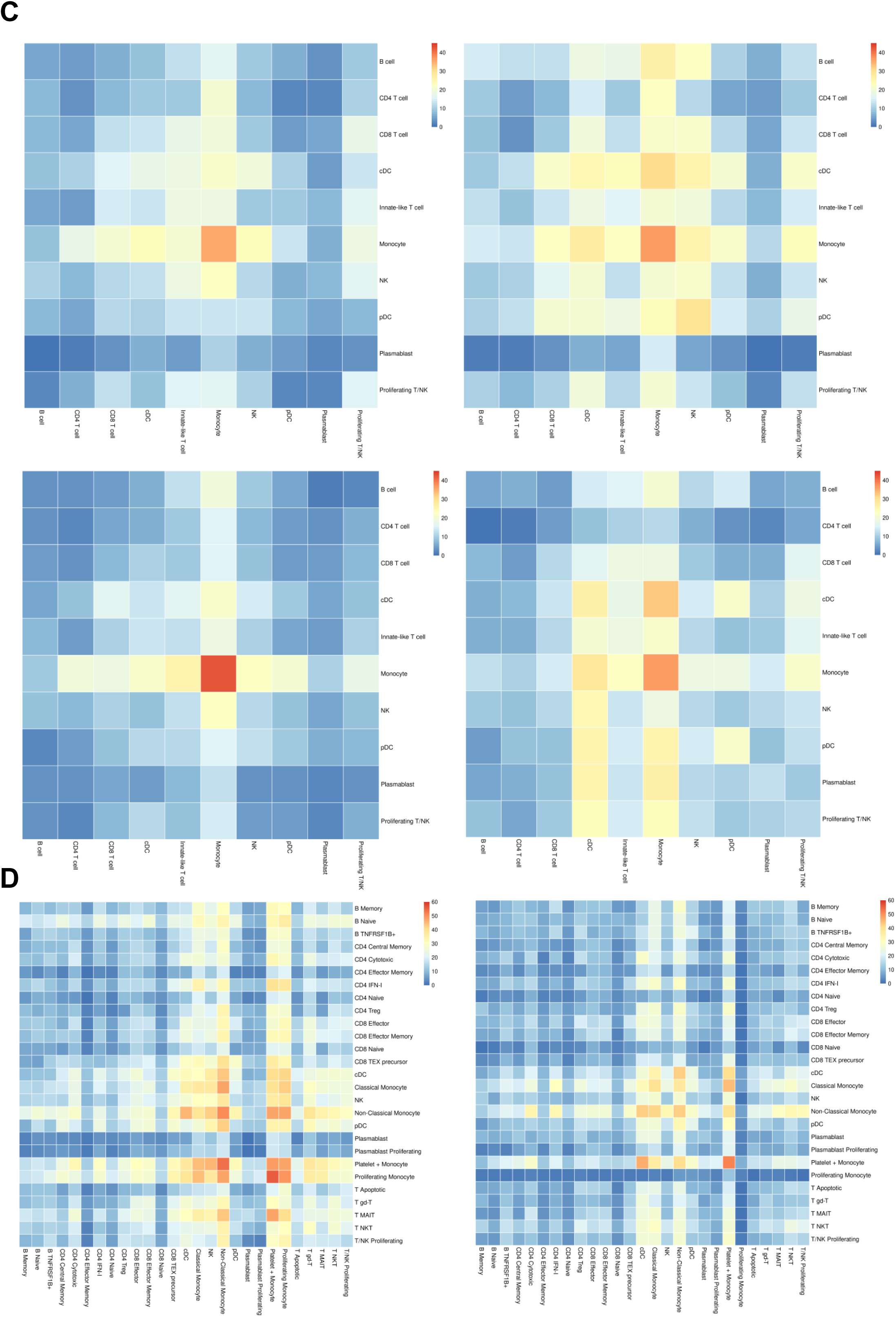
Comparative receptor-ligand interactions in COVID-19 and HIV-1^+^ patients. (A) Chord plots illustrating the top 25% expressed putative receptor-ligand interactions in COVID-19 (left) and HIV-1^+^ (right) patients between monocytes and DCs and adaptive immune cells. Interactions are depicted with the base of the arrow indicating the ligand, and the head of the arrow indicating the receptor. (B) As in (A), but split across nonventilated COVID-19 patients (top left), ventilated COVID-19 patients (top right), acute HIV-1^+^ patients (bottom left) and chronic HIV-1^+^ patients (bottom right). (C) Heatmap of the number of receptor-ligand interactions between each broad cell type in nonventilated COVID-19 patients (top left), ventilated COVID-19 patients (top right), acute HIV-1^+^ patients (bottom left) and chronic HIV-1^+^ patients (bottom right). (D) Heatmap of the number of receptor-ligand interactions between each cell type in COVID-19 patients (left), and HIV-1^+^ patients (right).

**Figure S6:**
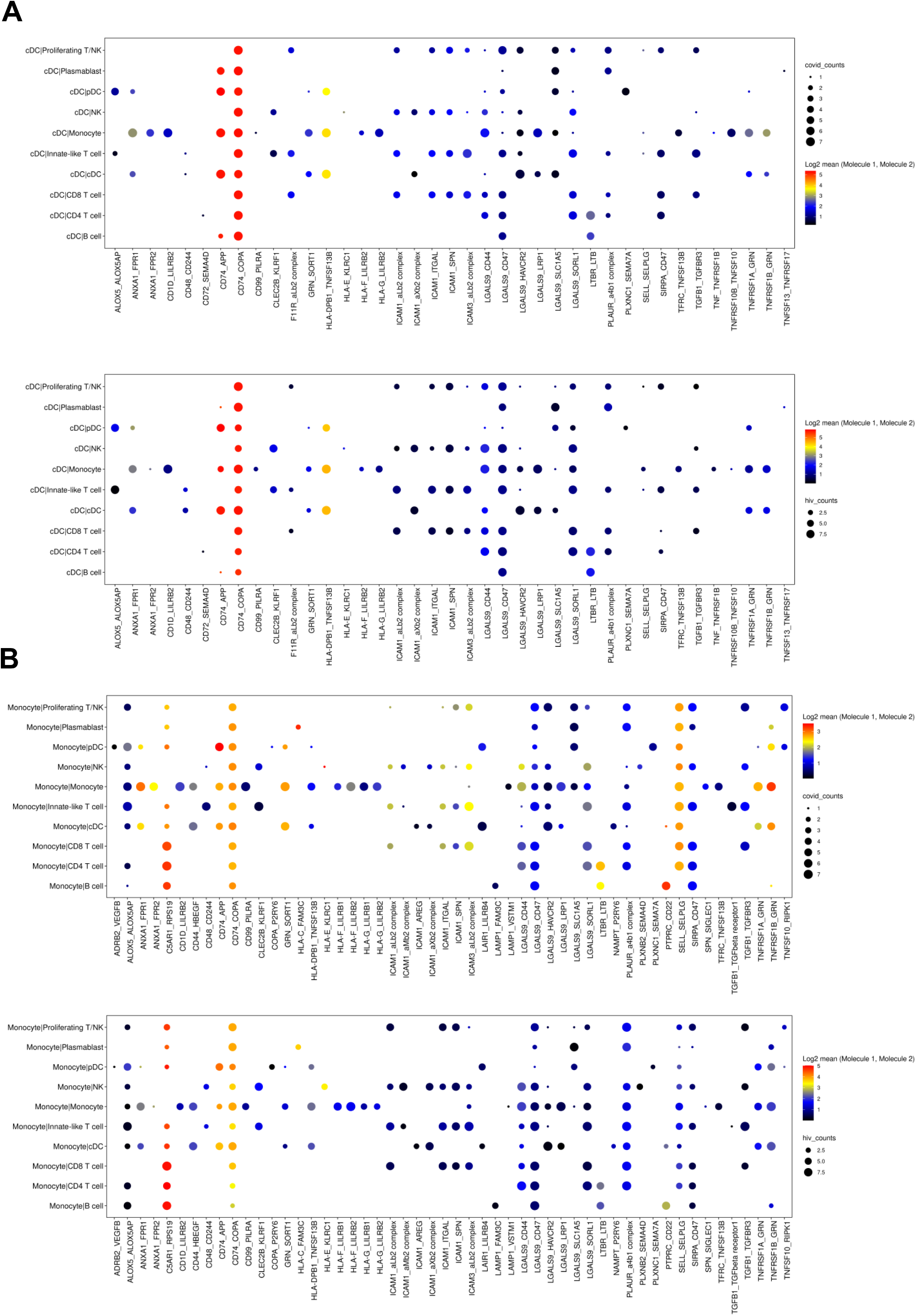
Receptor-ligand interactions of DCs and Monocytes. (A) Dot plot of receptor-ligand interactions between cDCs and other immune cells in COVID-19 patients (top) versus HIV-1^+^ patients (bottom). Color of each dot corresponds to the log2 of the mean expression of the interaction. Size of the dot corresponds to the number of patients the interaction was found to be significant in. (B) as in (A), but between monocytes and other immune cells.

**Figure S7:**
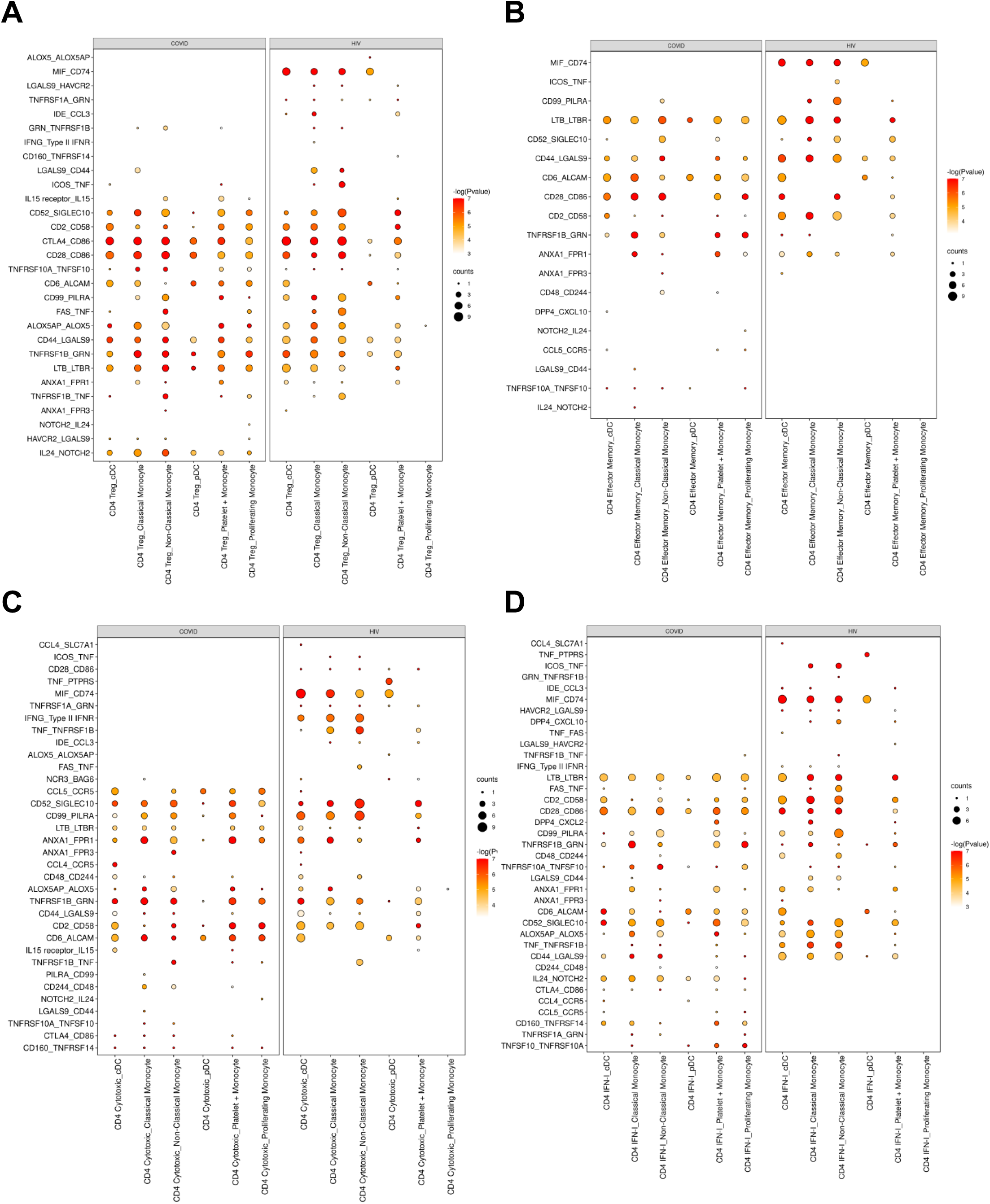
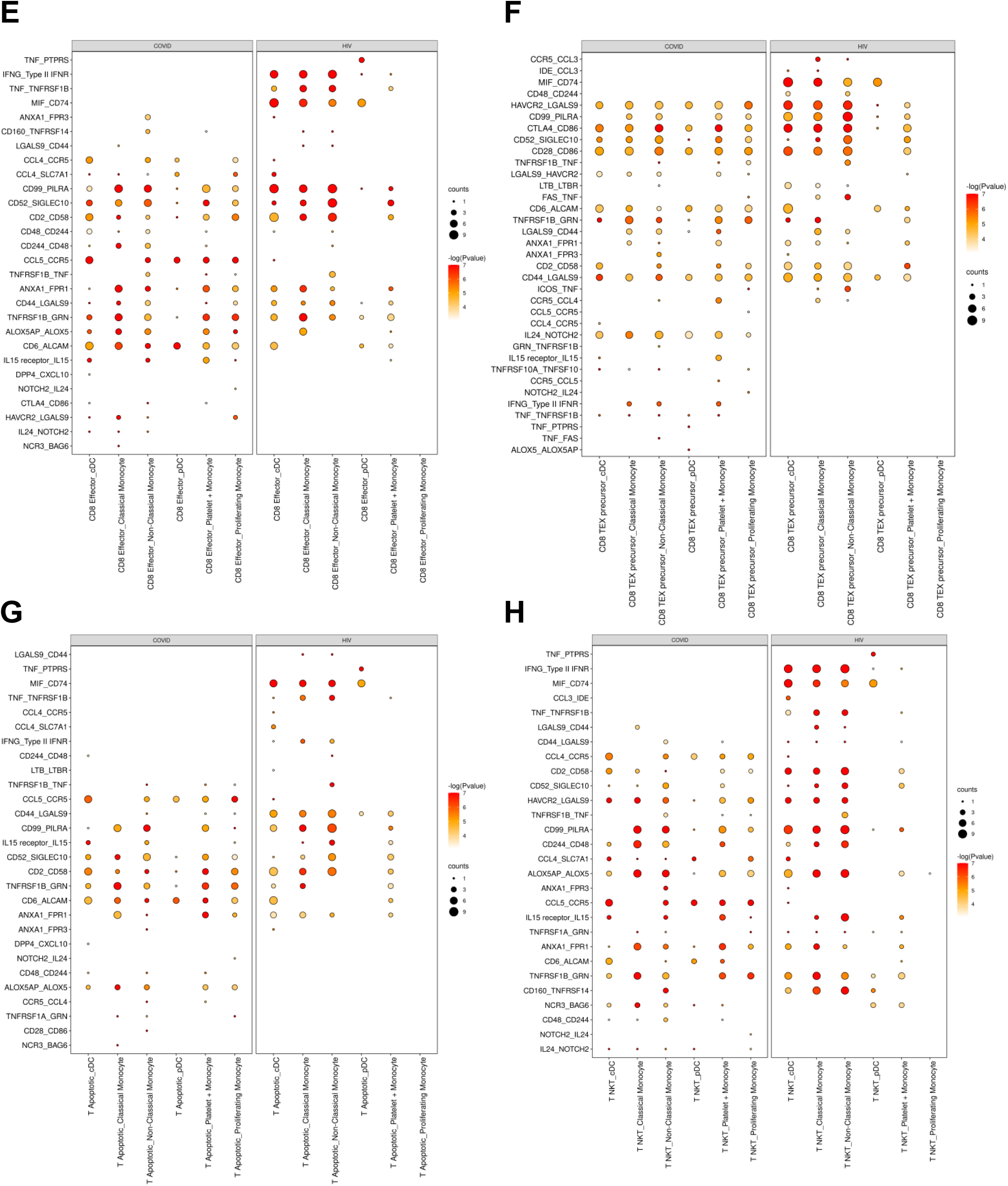
Receptor-ligand interactions between DCs/monocytes and CD4^+^/CD8^+^ T cells. (A) Dot plot of selected receptor-ligand interactions between Tregs and monocytes/DCs in COVID-19 patients (left) versus HIV-1^+^ patients (right). Color of each dot corresponds to the inverse log of the P value of the interaction. Size of the dot corresponds to the number of patients the interaction was found to be significant in. (B) As in (A), but with effector memory CD4^+^ T cells. (C) As in (A), but with cytotoxic CD4^+^ T cells. (D) As in (A), but with IFN-I^+^ CD4^+^ T cells. (E) As in (A), but with effector CD8^+^ T cells. (F) As in (A), but with precursor exhausted CD8^+^ T cells. (G) As in (A), but with apoptotic T cells. (H) As in (A), but with NKT cells

## References

10xGenomics (2020). PBMCs from a Healthy Donor: Whole Transcriptome Analysis. x. Genomics.

Abdelaal, T., Michielsen, L., Cats, D., Hoogduin, D., Mei, H., Reinders, M.J.T., and Mahfouz, A. (2019). A comparison of automatic cell identification methods for single-cell RNA sequencing data. Genome Biology 20, 194. 10.1186/s13059-019-1795-z.

Abedi, F., Hayes, A.W., Reiter, R., and Karimi, G. (2020a). Acute lung injury: The therapeutic role of Rho kinase inhibitors. Pharmacological Research 155, 104736. 10.1016/j.phrs.2020.104736.

Abedi, F., Rezaee, R., and Karimi, G. (2020b). Plausibility of therapeutic effects of Rho kinase inhibitors against Severe Acute Respiratory Syndrome Coronavirus 2 (COVID-19). Pharmacological Research 156, 104808. 10.1016/j.phrs.2020.104808.

Abraham, S., Choi, J.G., Ortega, N.M., Zhang, J.L., Shankar, P., and Swamy, N.M. (2016). Gene therapy with plasmids encoding IFN-beta or IFN-alpha 14 confers long-term resistance to HIV-1 in humanized mice. Oncotarget 7, 78412–78420. 10.18632/oncotarget.12512.

Akbay, B., Shmakova, A., Vassetzky, Y., and Dokudovskaya, S. (2020). Modulation of mTORC1 Signaling Pathway by HIV-1. Cells 9. 10.3390/cells9051090.

Ampudia, J., Young-Greenwald, W.W., Badrani, J., Gatto, S., Pavlicek, A., Doherty, T., Connelly, S., and Ng, C.T. (2020). CD6-ALCAM signaling regulates multiple effector/memory T cell functions. Journal of Immunology 204.

Aran, D., Looney, A.P., Liu, L., Wu, E., Fong, V., Hsu, A., Chak, S., Naikawadi, R.P., Wolters, P.J., Abate, A.R., et al. (2019). Reference-based analysis of lung single-cell sequencing reveals a transitional profibrotic macrophage. Nature Immunology 20, 163-+. 10.1038/s41590-018-0276-y.

Baden, L.R., El Sahly, H.M., Essink, B., Kotloff, K., Frey, S., Novak, R., Diemert, D., Spector, S.A., Rouphael, N., Creech, C.B., et al. (2021). Efficacy and Safety of the mRNA-1273 SARS-CoV-2 Vaccine. N Engl J Med 384, 403–416. 10.1056/NEJMoa2035389.

Baum, L.L. (2010). Role of humoral immunity in host defense against HIV. Curr HIV/AIDS Rep 7, 11–18. 10.1007/s11904-009-0036-6.

Bernardes, J.P., Mishra, N., Tran, F., Bahmer, T., Best, L., Blase, J.I., Bordoni, D., Franzenburg, J., Geisen, U., Josephs-Spaulding, J., et al. (2020). Longitudinal Multi-omics Analyses Identify Responses of Megakaryocytes, Erythroid Cells, and Plasmablasts as Hallmarks of Severe COVID-19. Immunity 53, 1296-+. 10.1016/j.immuni.2020.11.017.

Bieberich, F., Vazquez-Lombardi, R., Yermanos, A., Ehling, R.A., Mason, D.M., Wagner, B., Kapetanovic, E., Di Roberto, R.B., Weber, C.R., Savic, M., et al. (2021). A Single-Cell Atlas of Lymphocyte Adaptive Immune Repertoires and Transcriptomes Reveals Age-Related Differences in Convalescent COVID-19 Patients. Front Immunol 12, 701085. 10.3389/fimmu.2021.701085.

Botbol, Y., Guerrero-Ros, I., and Macian, F. (2016). Key roles of autophagy in regulating T-cell function. European Journal of Immunology 46, 1326–1334. 10.1002/eji.201545955.

Brenchley, J.M., Price, D.A., Schacker, T.W., Asher, T.E., Silvestri, G., Rao, S., Kazzaz, Z., Bornstein, E., Lambotte, O., Altmann, D., et al. (2006). Microbial translocation is a cause of systemic immune activation in chronic HIV infection. Nat Med 12, 1365–1371. 10.1038/nm1511.

Buggert, M., Nguyen, S., Salgado-Montes de Oca, G., Bengsch, B., Darko, S., Ransier, A., Roberts, E.R., Del Alcazar, D., Brody, I.B., Vella, L.A., et al. (2018). Identification and characterization of HIV-specific resident memory CD8. Sci Immunol 3. 10.1126/sciimmunol.aar4526.

Cai, Y., Liu, Y., and Zhang, X. (2007). Suppression of coronavirus replication by inhibition of the MEK signaling pathway. Journal of Virology 81, 446–456. 10.1128/jvi.01705-06.

Campbell, J.H., Hearps, A.C., Martin, G.E., Williams, K.C., and Crowe, S.M. (2014). The importance of monocytes and macrophages in HIV pathogenesis, treatment, and cure. Aids 28, 2175–2187. 10.1097/qad.0000000000000408.

Chen, G., Kroemer, G., and Kepp, O. (2020a). Mitophagy: An Emerging Role in Aging and Age-Associated Diseases. Frontiers in Cell and Developmental Biology 8, 200. 10.3389/fcell.2020.00200.

Chen, G., Ning, B., and Shi, T. (2019a). Single-Cell RNA-Seq Technologies and Related Computational Data Analysis. Frontiers in Genetics 10, 317. 10.3389/fgene.2019.00317.

Chen, G., Wu, D., Guo, W., Cao, Y., Huang, D., Wang, H., Wang, T., Zhang, X., Chen, H., Yu, H., et al. (2020b). Clinical and immunological features of severe and moderate coronavirus disease 2019. Journal of Clinical Investigation 130, 2620–2629. 10.1172/jci137244.

Chen, J.-X., Xu, X., and Zhang, S. (2019b). Silence of long noncoding RNA NEAT1 exerts suppressive effects on immunity during sepsis by promoting microRNA-125-dependent MCEMP1 downregulation. Iubmb Life 71, 956–968. 10.1002/iub.2033.

Choi, U.Y., Kang, J.S., Hwang, Y.S., and Kim, Y.J. (2015). Oligoadenylate synthase-like (OASL) proteins: dual functions and associations with diseases. Exp Mol Med 47, e144. 10.1038/emm.2014.110.

Chow, R.D., Majety, M., and Chen, S. (2021). The aging transcriptome and cellular landscape of the human lung in relation to SARS-CoV-2. Nat Commun 12, 4. 10.1038/s41467-020-20323-9.

Coleman, C.M., and Wu, L. (2009). HIV interactions with monocytes and dendritic cells: viral latency and reservoirs. Retrovirology 6, 51. 10.1186/1742-4690-6-51.

Costela-Ruiz, V.J., Illescas-Montes, R., Puerta-Puerta, J.M., Ruiz, C., and Melguizo-Rodriguez, L. (2020). SARS-CoV-2 infection: The role of cytokines in COVID-19 disease. Cytokine & Growth Factor Reviews 54, 62–75. 10.1016/j.cytogfr.2020.06.001.

Cotugno, N., Ruggiero, A., Bonfante, F., Petrara, M.R., Zicari, S., Pascucci, G.R., Zangari, P., De Ioris, M.A., Santilli, V., Manno, E.C., et al. (2021). Virological and immunological features of SARS-CoV-2- infected children who develop neutralizing antibodies. Cell Rep 34, 108852. 10.1016/j.celrep.2021.108852.

Dangi, T., Class, J., Palacio, N., Richner, J.M., and Penaloza MacMaster, P. (2021a). Combining spike- and nucleocapsid-based vaccines improves distal control of SARS-CoV-2. Cell Rep 36, 109664. 10.1016/j.celrep.2021.109664.

Dangi, T., Palacio, N., Sanchez, S., Park, M., Class, J., Visvabharathy, L., Ciucci, T., Koralnik, I.J., Richner, J.M., and Penaloza-MacMaster, P. (2021b). Cross-protective immunity following coronavirus vaccination and coronavirus infection. J Clin Invest 131. 10.1172/JCI151969.

De Biasi, S., Lo Tartaro, D., Meschiari, M., Gibellini, L., Bellinazzi, C., Borella, R., Fidanza, L., Mattioli, M., Paolini, A., Gozzi, L., et al. (2020). Expansion of plasmablasts and loss of memory B cells in peripheral blood from COVID-19 patients with pneumonia. European Journal of Immunology 50, 1283–1294. 10.1002/eji.202048838.

de la Rica, R., Borges, M., and Gonzalez-Freire, M. (2020). COVID-19: In the Eye of the Cytokine Storm. Frontiers in Immunology 11, 558898. 10.3389/fimmu.2020.558898.

Deeks, S.G., Tracy, R., and Douek, D.C. (2013). Systemic effects of inflammation on health during chronic HIV infection. Immunity 39, 633–645. 10.1016/j.immuni.2013.10.001.

Delorey, T.M., Ziegler, C.G.K., Heimberg, G., Normand, R., Yang, Y., Segerstolpe, A., Abbondanza, D., Fleming, S.J., Subramanian, A., Montoro, D.T., et al. (2021). COVID-19 tissue atlases reveal SARS-CoV- 2 pathology and cellular targets. Nature 595, 107-+. 10.1038/s41586-021-03570-8.

Doitsh, G., Galloway, N.L.K., Geng, X., Yang, Z., Monroe, K.M., Zepeda, O., Hunt, P.W., Hatano, H., Sowinski, S., Munoz-Arias, I., and Greene, W.C. (2014). Cell death by pyroptosis drives CD4 T-cell depletion in HIV-1 infection. Nature 505, 509-+. 10.1038/nature12940.

Dong, E., Du, H., and Gardner, L. (2020). An interactive web-based dashboard to track COVID-19 in real time. Lancet Infect Dis 20, 533–534. 10.1016/S1473-3099(20)30120-1.

Efremova, M., Vento-Tormo, M., Teichmann, S.A., and Vento-Tormo, R. (2020). CellPhoneDB: inferring cell-cell communication from combined expression of multi-subunit ligand-receptor complexes. Nat Protoc 15, 1484–1506. 10.1038/s41596-020-0292-x.

Fergusson, J.R., Hühn, M.H., Swadling, L., Walker, L.J., Kurioka, A., Llibre, A., Bertoletti, A., Holländer, G., Newell, E.W., Davis, M.M., et al. (2016). CD161(int)CD8+ T cells: a novel population of highly functional, memory CD8+ T cells enriched within the gut. Mucosal Immunol 9, 401–413. 10.1038/mi.2015.69.

Galani, I.E., Rovina, N., Lampropoulou, V., Triantafyllia, V., Manioudaki, M., Pavlos, E., Koukaki, E., Fragkou, P.C., Panou, V., Rapti, V., et al. (2021). Untuned antiviral immunity in COVID-19 revealed by temporal type I/III interferon patterns and flu comparison. Nat Immunol 22, 32–40. 10.1038/s41590-020-00840-x.

Ganji, R., and Reddy, P.H. (2021). Impact of COVID-19 on Mitochondrial-Based Immunity in Aging and Age-Related Diseases. Frontiers in Aging Neuroscience 12, 614650. 10.3389/fnagi.2020.614650.

Garg, H., Mohl, J., and Joshi, A. (2012). HIV-1 Induced Bystander Apoptosis. Viruses-Basel 4, 3020–3043. 10.3390/v4113020.

Gassen, N.C., Papies, J., Bajaj, T., Emanuel, J., Dethloff, F., Chua, R.L., Trimpert, J., Heinemann, N., Niemeyer, C., Weege, F., et al. (2021). SARS-CoV-2-mediated dysregulation of metabolism and autophagy uncovers host-targeting antivirals. Nature Communications 12, 3818. 10.1038/s41467-021-24007-w.

Gasteiger, G., and Rudensky, A.Y. (2014). Interactions between innate and adaptive lymphocytes. Nat Rev Immunol 14, 631–639. 10.1038/nri3726.

Gawad, C., Koh, W., and Quake, S.R. (2016). Single-cell genome sequencing: current state of the science. Nature Reviews Genetics 17, 175–188. 10.1038/nrg.2015.16.

Goel, R.R., Painter, M.M., Apostolidis, S.A., Mathew, D., Meng, W., Rosenfeld, A.M., Lundgreen, K.A., Reynaldi, A., Khoury, D.S., Pattekar, A., et al. (2021a). mRNA vaccines induce durable immune memory to SARS-CoV-2 and variants of concern. Science 374, abm0829. 10.1126/science.abm0829.

Goel, S., Saheb Sharif-Askari, F., Saheb Sharif Askari, N., Madkhana, B., Alwaa, A.M., Mahboub, B., Zakeri, A.M., Ratemi, E., Hamoudi, R., Hamid, Q., and Halwani, R. (2021b). SARS-CoV-2 Switches ’on’ MAPK and NF kappa B Signaling via the Reduction of Nuclear DUSP1 and DUSP5 Expression. Frontiers in Pharmacology 12, 631879. 10.3389/fphar.2021.631879.

Goveia, J., Rohlenova, K., Taverna, F., Treps, L., Conradi, L.C., Pircher, A., Geldhof, V., de Rooij, L.P.M.H., Kalucka, J., Sokol, L., et al. (2020). An Integrated Gene Expression Landscape Profiling Approach to Identify Lung Tumor Endothelial Cell Heterogeneity and Angiogenic Candidates. Cancer Cell 37, 421. 10.1016/j.ccell.2020.03.002.

Guarda, G., Braun, M., Staehli, F., Tardivel, A., Mattmann, C., Foerster, I., Farlik, M., Decker, T., Du Pasquier, R.A., Romero, P., and Tschopp, J. (2011). Type I Interferon Inhibits Interleukin-1 Production and Inflammasome Activation. Immunity 34, 213–223. 10.1016/j.immuni.2011.02.006.

Hao, Y., Hao, S., Andersen-Nissen, E., Mauck, W.M.I.I.I.I.I.I., Zheng, S., Butler, A., Lee, M.J., Wilk, A.J., Darby, C., Zager, M., et al. (2021). Integrated analysis of multimodal single-cell data. Cell 184, 3573-+. 10.1016/j.cell.2021.04.048.

Haque, A., Engel, J., Teichmann, S.A., and Lonnberg, T. (2017). A practical guide to single-cell RNA-sequencing for biomedical research and clinical applications. Genome Medicine 9, 75. 10.1186/s13073-017-0467-4.

Hasan, M.Z., Islam, S., Matsumoto, K., and Kawai, T. (2021). SARS-CoV-2 infection initiates interleukin-17-enriched transcriptional response in different cells from multiple organs. Sci Rep 11, 16814. 10.1038/s41598-021-96110-3.

Hodge, R.G., and Ridley, A.J. (2016). Regulating Rho GTPases and their regulators. Nature Reviews Molecular Cell Biology 17, 496–510. 10.1038/nrm.2016.67.

Hoe, E., McKay, F., Schibeci, S., Heard, R., Stewart, G., and Booth, D. (2010). Interleukin 7 Receptor Alpha Chain Haplotypes Vary in Their Influence on Multiple Sclerosis Susceptibility and Response to Interferon Beta. Journal of Interferon and Cytokine Research 30, 291–298. 10.1089/jir.2009.0060.

Hoffmann, M., Kleine-Weber, H., Schroeder, S., Kruger, N., Herrler, T., Erichsen, S., Schiergens, T.S., Herrler, G., Wu, N.H., Nitsche, A., et al. (2020). SARS-CoV-2 Cell Entry Depends on ACE2 and TMPRSS2 and Is Blocked by a Clinically Proven Protease Inhibitor. Cell 181, 271-+. 10.1016/j.cell.2020.02.052.

Hu, H., Juvekar, A., Lyssiotis, C.A., Lien, E.C., Albeck, J.G., Oh, D., Varma, G., Hung, Y.P., Ullas, S., Lauring, J., et al. (2016). Phosphoinositide 3-Kinase Regulates Glycolysis through Mobilization of Aldolase from the Actin Cytoskeleton. Cell 164, 433–446. 10.1016/j.cell.2015.12.042.

Huang, Q., Liu, Y., Du, Y., and Garmire, L.X. (2021). Evaluation of Cell Type Annotation R Packages on Single-cell RNA-seq Data. Genomics Proteomics & Bioinformatics 19, 267–281. 10.1016/j.gpb.2020.07.004.

Hui, E., Cheung, J., Zhu, J., Su, X., Taylor, M.J., Wallweber, H.A., Sasmal, D.K., Huang, J., Kim, J.M., Mellman, I., and Vale, R.D. (2017). T cell costimulatory receptor CD28 is a primary target for PD-1-mediated inhibition. Science 355, 1428–1433. 10.1126/science.aaf1292.

Hömig-Hölzel, C., Hojer, C., Rastelli, J., Casola, S., Strobl, L.J., Müller, W., Quintanilla-Martinez, L., Gewies, A., Ruland, J., Rajewsky, K., and Zimber-Strobl, U. (2008). Constitutive CD40 signaling in B cells selectively activates the noncanonical NF-kappaB pathway and promotes lymphomagenesis. J Exp Med 205, 1317–1329. 10.1084/jem.20080238.

Jha, P., and Das, H. (2017). KLF2 in Regulation of NF-κB-Mediated Immune Cell Function and Inflammation. Int J Mol Sci 18. 10.3390/ijms18112383.

Jin, X., Zhou, W., Luo, M., Wang, P., Xu, Z., Ma, K., Cao, H., Xu, C., Huang, Y., Cheng, R., et al. (2021). Global characterization of B cell receptor repertoire in COVID-19 patients by single-cell V(D)J sequencing. Brief Bioinform 22. 10.1093/bib/bbab192.

Jones, S.A., and Hunter, C.A. (2021). Is IL-6 a key cytokine target for therapy in COVID-19? Nature Reviews Immunology 21, 337–339. 10.1038/s41577-021-00553-8.

Kasuga, Y., Zhu, B., Jang, K.-J., and Yoo, J.-S. (2021). Innate immune sensing of coronavirus and viral evasion strategies. Experimental and Molecular Medicine 53, 723–736. 10.1038/s12276-021-00602-1.

Kazer, S.W., Aicher, T.P., Muema, D.M., Carroll, S.L., Ordovas-Montanes, J., Miao, V.N., Tu, A.A., Ziegler, C.G.K., Nyquist, S.K., Wong, E.B., et al. (2020). Integrated single-cell analysis of multicellular immune dynamics during hyperacute HIV-1 infection. Nature Medicine 26, 511-+. 10.1038/s41591-020-0799-2.

Kedzierska, K., and Crowe, S.M. (2002). The role of monocytes and macrophages in the pathogenesis of HIV-1 infection. Curr Med Chem 9, 1893–1903. 10.2174/0929867023368935.

Khan, M., Syed, G.H., Kim, S.J., and Siddiqui, A. (2015). Mitochondrial dynamics and viral infections: A close nexus. Biochim Biophys Acta 1853, 2822–2833. 10.1016/j.bbamcr.2014.12.040.

Konduri, V., Oyewole-Said, D., Vazquez-Perez, J., Weldon, S.A., Halpert, M.M., Levitt, J.M., and Decker, W.K. (2020). CD8. Front Immunol 11, 613204. 10.3389/fimmu.2020.613204.

Kono, M., Tatsumi, K., Imai, A.M., Saito, K., Kuriyama, T., and Shirasawa, H. (2008). Inhibition of human coronavirus 229E infection in human epithelial lung cells (L132) by chloroquine: Involvement of p38 MAPK and ERK. Antiviral Research 77, 150–152. 10.1016/j.antiviral.2007.10.011.

Kopitar-Jerala, N. (2017). The Role of Interferons in Inflammation and Inflammasome Activation. Front Immunol 8, 873. 10.3389/fimmu.2017.00873.

Korsunsky, I., Millard, N., Fan, J., Slowikowski, K., Zhang, F., Wei, K., Baglaenko, Y., Brenner, M., Loh, P.R., and Raychaudhuri, S. (2019). Fast, sensitive and accurate integration of single-cell data with Harmony. Nat Methods 16, 1289–1296. 10.1038/s41592-019-0619-0.

Kovacs, J.R., Li, C., Yang, Q., Li, G., Garcia, I.G., Ju, S., Roodman, D.G., Windle, J.J., Zhang, X., and Lu, B. (2012). Autophagy promotes T-cell survival through degradation of proteins of the cell death machinery. Cell Death and Differentiation 19, 144–152. 10.1038/cdd.2011.78.

Krzak, M., Raykov, Y., Boukouvalas, A., Cutillo, L., and Angelini, C. (2019). Benchmark and Parameter Sensitivity Analysis of Single-Cell RNA Sequencing Clustering Methods. Frontiers in Genetics 10, 1253. 10.3389/fgene.2019.01253.

Kuksin, M., Morel, D., Aglave, M., Danlos, F.-X., Marabelle, A., Zinovyev, A., Gautheret, D., and Verlingue, L. (2021). Applications of single-cell and bulk RNA sequencing in onco-immunology. European Journal of Cancer 149, 193–210. 10.1016/j.ejca.2021.03.005.

Kuri-Cervantes, L., Pampena, M.B., Meng, W., Rosenfeld, A.M., Ittner, C.A.G., Weisman, A.R., Agyekum, R.S., Mathew, D., Baxter, A.E., Vella, L.A., et al. (2020). Comprehensive mapping of immune perturbations associated with severe COVID-19. Science Immunology 5, eabd7114. 10.1126/sciimmunol.abd7114.

Langfelder, P., and Horvath, S. (2008). WGCNA: an R package for weighted correlation network analysis. BMC Bioinformatics 9, 559. 10.1186/1471-2105-9-559.

Lavender, K.J., Gibbert, K., Peterson, K.E., Van Dis, E., Francois, S., Woods, T., Messer, R.J., Gawanbacht, A., Müller, J.A., Münch, J., et al. (2016). Interferon Alpha Subtype-Specific Suppression of HIV-1 Infection In Vivo. J Virol 90, 6001–6013. 10.1128/JVI.00451-16.

Lee, J.S., Park, S., Jeong, H.W., Ahn, J.Y., Choi, S.J., Lee, H., Choi, B., Nam, S.K., Sa, M., Kwon, J.-S., et al. (2020). Immunophenotyping of COVID-19 and influenza highlights the role of type I interferons in development of severe COVID-19. Science Immunology 5, eabd1554. 10.1126/sciimmunol.abd1554.

Lee, J.S., and Shin, E.-C. (2020). The type I interferon response in COVID-19: implications for treatment. Nature Reviews Immunology 20, 585–586. 10.1038/s41577-020-00429-3.

Lee, J.W., Su, Y., Baloni, P., Chen, D., Pavlovitch-Bedzyk, A.J., Yuan, D., Duvvuri, V.R., Ng, R.H., Choi, J., Xie, J., et al. (2021). Integrated analysis of plasma and single immune cells uncovers metabolic changes in individuals with COVID-19. Nature Biotechnology. 10.1038/s41587-021-01020-4.

Li, S. (2019). Regulation of Ribosomal Proteins on Viral Infection. Cells 8. 10.3390/cells8050508.

Liao, M., Liu, Y., Yuan, J., Wen, Y., Xu, G., Zhao, J., Cheng, L., Li, J., Wang, X., Wang, F., et al. (2020). Single-cell landscape of bronchoalveolar immune cells in patients with COVID-19. Nature Medicine 26, 842-+. 10.1038/s41591-020-0901-9.

Liu, B., Li, C., Li, Z., Wang, D., Ren, X., and Zhang, Z. (2020). An entropy-based metric for assessing the purity of single cell populations. Nat Commun 11, 3155. 10.1038/s41467-020-16904-3.

Liu, N., Jiang, C., Cai, P., Shen, Z., Sun, W., Xu, H., Fang, M., Yao, X., Zhu, L., Gao, X., et al. (2021). Single-cell analysis of COVID-19, sepsis, and HIV infection reveals hyperinflammatory and immunosuppressive signatures in monocytes. Cell Reports 37, 109793. 10.1016/j.celrep.2021.109793.

Locci, M., Havenar-Daughton, C., Landais, E., Wu, J., Kroenke, M.A., Arlehamn, C.L., Su, L.F., Cubas, R., Davis, M.M., Sette, A., et al. (2013). Human circulating PD-1+CXCR3-CXCR5+ memory Tfh cells are highly functional and correlate with broadly neutralizing HIV antibody responses. Immunity 39, 758–769. 10.1016/j.immuni.2013.08.031.

Longhitano, L., Tibullo, D., Giallongo, C., Lazzarino, G., Tartaglia, N., Galimberti, S., Li Volti, G., Palumbo, G.A., and Liso, A. (2020). Proteasome Inhibitors as a Possible Therapy for SARS-CoV-2. International Journal of Molecular Sciences 21. 10.3390/ijms21103622.

Lu, J., Pan, Q., Rong, L., Liu, S.-L., and Liang, C. (2011). The IFITM Proteins Inhibit HIV-1 Infection. Journal of Virology 85, 2126–2137. 10.1128/jvi.01531-10.

Lu, T.T., and Browning, J.L. (2014). Role of the Lymphotoxin/LIGHT System in the Development and Maintenance of Reticular Networks and Vasculature in Lymphoid Tissues. Front Immunol 5, 47. 10.3389/fimmu.2014.00047.

Ma, A., Zhang, L., Ye, X., Chen, J., Yu, J., Zhuang, L., Weng, C., Petersen, F., Wang, Z., and Yu, X. (2021). High Levels of Circulating IL-8 and Soluble IL-2R Are Associated With Prolonged Illness in Patients With Severe COVID-19. Frontiers in Immunology 12, 626235. 10.3389/fimmu.2021.626235.

Macedo, A.B., Novis, C.L., and Bosque, A. (2019). Targeting Cellular and Tissue HIV Reservoirs With Toll-Like Receptor Agonists. Front Immunol 10, 2450. 10.3389/fimmu.2019.02450.

MacParland, S.A., Liu, J.C., Ma, X.Z., Innes, B.T., Bartczak, A.M., Gage, B.K., Manuel, J., Khuu, N., Echeverri, J., Linares, I., et al. (2018). Single cell RNA sequencing of human liver reveals distinct intrahepatic macrophage populations. Nature Communications 9, 4383. 10.1038/s41467-018-06318-7.

Malarkannan, S. (2020). NKG7 makes a better killer. Nature Immunology 21, 1139–1140. 10.1038/s41590-020-0767-5.

Manches, O., Frleta, D., and Bhardwaj, N. (2014). Dendritic cells in progression and pathology of HIV infection. Trends in Immunology 35, 114–122. 10.1016/j.it.2013.10.003.

Mashayekhi-Sardoo, H., and Hosseinjani, H. (2021). A new application of mTOR inhibitor drugs as potential therapeutic agents for COVID-19. J Basic Clin Physiol Pharmacol. 10.1515/jbcpp-2020-0495.

Mayer-Barber, K.D., and Yan, B. (2017). Clash of the Cytokine Titans: counter-regulation of interleukin-1 and type I interferon-mediated inflammatory responses. Cellular & Molecular Immunology 14, 22–35. 10.1038/cmi.2016.25.

McNab, F., Mayer-Barber, K., Sher, A., Wack, A., and O’Garra, A. (2015). Type I interferons in infectious disease. Nature Reviews Immunology 15, 87–103. 10.1038/nri3787.

Meier, A., and Altfeld, M. (2007). Toll-like receptor signaling in HIV-1 infection: a potential target for therapy? Expert Review of Anti-Infective Therapy 5, 323–326. 10.1586/14787210.5.3.323.

Melms, J.C., Biermann, J., Huang, H., Wang, Y., Nair, A., Tagore, S., Katsyv, I., Rendeiro, A.F., Amin, A.D., Schapiro, D., et al. (2021). A molecular single-cell lung atlas of lethal COVID-19. Nature 595, 114-+. 10.1038/s41586-021-03569-1.

Mercado, N.B., Zahn, R., Wegmann, F., Loos, C., Chandrashekar, A., Yu, J., Liu, J., Peter, L., McMahan, K., Tostanoski, L.H., et al. (2020). Single-shot Ad26 vaccine protects against SARS-CoV-2 in rhesus macaques. Nature 586, 583–588. 10.1038/s41586-020-2607-z.

Meås, H.Z., Haug, M., Beckwith, M.S., Louet, C., Ryan, L., Hu, Z., Landskron, J., Nordbø, S.A., Taskén, K., Yin, H., et al. (2020). Sensing of HIV-1 by TLR8 activates human T cells and reverses latency. Nat Commun 11, 147. 10.1038/s41467-019-13837-4.

Mizutani, T., Fukushi, S., Saijo, M., Kurane, I., and Morikawa, S. (2005). JNK and PI3k/Akt signaling pathways are required for establishing persistent SARS-CoV infection in Vero E6 cells. Biochim Biophys Acta 1741, 4–10. 10.1016/j.bbadis.2005.04.004.

Moir, S., and Fauci, A.S. (2009). B cells in HIV infection and disease. Nature Reviews Immunology 9, 235–245. 10.1038/nri2524.

Monaco, G., Lee, B., Xu, W., Mustafah, S., Hwang, Y.Y., Carré, C., Burdin, N., Visan, L., Ceccarelli, M., Poidinger, M., et al. (2019). RNA-Seq Signatures Normalized by mRNA Abundance Allow Absolute Deconvolution of Human Immune Cell Types. Cell Rep 26, 1627–1640.e1627. 10.1016/j.celrep.2019.01.041.

Moseman, E.A., Wu, T., de la Torre, J.C., Schwartzberg, P.L., and McGavern, D.B. (2016). Type I interferon suppresses virus-specific B cell responses by modulating CD8. Sci Immunol 1. 10.1126/sciimmunol.aah3565.

Mutvei, A.P., Nagiec, M.J., Hamann, J.C., Kim, S.G., Vincent, C.T., and Blenis, J. (2020). Rap1-GTPases control mTORC1 activity by coordinating lysosome organization with amino acid availability. Nat Commun 11, 1416. 10.1038/s41467-020-15156-5.

Nguyen, T.H., McAuley, J.L., Kim, Y., Zheng, M.Z., Gherardin, N.A., Godfrey, D.I., Purcell, D.F., Sullivan, L.C., Westall, G.P., Reading, P.C., et al. (2021). Influenza, but not SARS-CoV-2, infection induces a rapid interferon response that wanes with age and diminished tissue-resident memory CD8. Clin Transl Immunology 10, e1242. 10.1002/cti2.1242.

Okhotnikov, K., Charpentier, T., and Cadars, S. (2016). Supercell program: a combinatorial structure-generation approach for the local-level modeling of atomic substitutions and partial occupancies in crystals. J Cheminform 8, 17. 10.1186/s13321-016-0129-3.

Ouyang, W., and O’Garra, A. (2019). IL-10 Family Cytokines IL-10 and IL-22: from Basic Science to Clinical Translation. Immunity 50, 871–891. 10.1016/j.immuni.2019.03.020.

Pairo-Castineira, E., Clohisey, S., Klaric, L., Bretherick, A.D., Rawlik, K., Pasko, D., Walker, S., Parkinson, N., Fourman, M.H., Russell, C.D., et al. (2021). Genetic mechanisms of critical illness in COVID-19. Nature 591, 92–98. 10.1038/s41586-020-03065-y.

Palacio, N., Dangi, T., Chung, Y.R., Wang, Y., Loredo-Varela, J.L., Zhang, Z., and Penaloza-MacMaster, P. (2020). Early type I IFN blockade improves the efficacy of viral vaccines. J Exp Med 217. 10.1084/jem.20191220.

Pasquini, G., Arias, J.E.R., Schafer, P., and Busskamp, V. (2021). Automated methods for cell type annotation on scRNA-seq data. Computational and Structural Biotechnology Journal 19, 961–969. 10.1016/j.csbj.2021.01.015.

Pavlovic, M., Gross, C., Chili, C., Secher, T., and Treiner, E. (2020). MAIT Cells Display a Specific Response to Type 1 IFN Underlying the Adjuvant Effect of TLR7/8 Ligands. Front Immunol 11, 2097. 10.3389/fimmu.2020.02097.

PBMCs from a Healthy Donor: Whole Transcriptome Analysis. (2020). x. Genomics.

Peng, X., Ouyang, J., Isnard, S., Lin, J., Fombuena, B., Zhu, B., and Routy, J.P. (2020). Sharing CD4+ T Cell Loss: When COVID-19 and HIV Collide on Immune System. Front Immunol 11, 596631. 10.3389/fimmu.2020.596631.

Polack, F.P., Thomas, S.J., Kitchin, N., Absalon, J., Gurtman, A., Lockhart, S., Perez, J.L., Pérez Marc, G., Moreira, E.D., Zerbini, C., et al. (2020). Safety and Efficacy of the BNT162b2 mRNA Covid-19 Vaccine. N Engl J Med 383, 2603–2615. 10.1056/NEJMoa2034577.

Pua, H.H., Guo, J., Komatsu, M., and He, Y.-W. (2009). Autophagy Is Essential for Mitochondrial Clearance in Mature T Lymphocytes. Journal of Immunology 182, 4046–4055. 10.4049/jimmunol.0801143.

Regis, E.G., Barreto-de-Souza, V., Morgado, M.G., Bozza, M.T., Leng, L., Bucala, R., and Bou-Habib, D.C. (2010). Elevated levels of macrophage migration inhibitory factor (MIF) in the plasma of HIV-1-infected patients and in HIV-1-infected cell cultures: A relevant role on viral replication. Virology 399, 31–38. 10.1016/j.virol.2009.12.018.

Ren, X., Wen, W., Fan, X., Hou, W., Su, B., Cai, P., Li, J., Liu, Y., Tang, F., Zhang, F., et al. (2021). COVID-19 immune features revealed by a large-scale single-cell transcriptome atlas. Cell 184, 5838. 10.1016/j.cell.2021.10.023.

Reyfman, P.A., Walter, J.M., Joshi, N., Anekalla, K.R., McQuattie-Pimentel, A.C., Chiu, S., Fernandez, R., Akbarpour, M., Chen, C.I., Ren, Z.Y., et al. (2019). Single-Cell Transcriptomic Analysis of Human Lung Provides Insights into the Pathobiology of Pulmonary Fibrosis. American Journal of Respiratory and Critical Care Medicine 199, 1517–1536. 10.1164/rccm.201712-2410OC.

Roff, S.R., Noon-Song, E.N., and Yamamoto, J.K. (2014). The Significance of Interferon-γ in HIV-1 Pathogenesis, Therapy, and Prophylaxis. Front Immunol 4, 498. 10.3389/fimmu.2013.00498.

Rubin, E.J., Longo, D.L., and Baden, L.R. (2021). Interleukin-6 Receptor Inhibition in Covid-19-Cooling the Inflammatory Soup. New England Journal of Medicine 384, 1564–1565. 10.1056/NEJMe2103108.

Sanchez, S., Palacio, N., Dangi, T., Ciucci, T., and Penaloza-MacMaster, P. (2021). Fractionating a COVID-19 Ad5-vectored vaccine improves virus-specific immunity. Sci Immunol 6, eabi8635. 10.1126/sciimmunol.abi8635.

Sandler, N.G., Bosinger, S.E., Estes, J.D., Zhu, R.T., Tharp, G.K., Boritz, E., Levin, D., Wijeyesinghe, S., Makamdop, K.N., del Prete, G.Q., et al. (2014). Type I interferon responses in rhesus macaques prevent SIV infection and slow disease progression. Nature 511, 601–605. 10.1038/nature13554.

Satarker, S., Tom, A.A., Shaji, R.A., Alosious, A., Luvis, M., and Nampoothiri, M. (2021). JAK-STAT Pathway Inhibition and their Implications in COVID-19 Therapy. Postgraduate Medicine 133, 489–507. 10.1080/00325481.2020.1855921.

Schreiber, G. (2020). The Role of Type I Interferons in the Pathogenesis and Treatment of COVID-19. Frontiers in Immunology 11, 595739. 10.3389/fimmu.2020.595739.

Schultze, J.L., and Aschenbrenner, A.C. (2021). COVID-19 and the human innate immune system. Cell 184, 1671–1692. 10.1016/j.cell.2021.02.029.

Senoo, H., Kamimura, Y., Kimura, R., Nakajima, A., Sawai, S., Sesaki, H., and Iijima, M. (2019). Phosphorylated Rho-GDP directly activates mTORC2 kinase towards AKT through dimerization with Ras-GTP to regulate cell migration. Nat Cell Biol 21, 867–878. 10.1038/s41556-019-0348-8.

Singh, K.K., Chaubey, G., Chen, J.Y., and Suravajhala, P. (2020). Decoding SARS-CoV-2 hijacking of host mitochondria in COVID-19 pathogenesis. American Journal of Physiology-Cell Physiology 319, C258–C267. 10.1152/ajpcell.00224.2020.

Soper, A., Kimura, I., Nagaoka, S., Konno, Y., Yamamoto, K., Koyanagi, Y., and Sato, K. (2017). Type I Interferon Responses by HIV-1 Infection: Association with Disease Progression and Control. Front Immunol 8, 1823. 10.3389/fimmu.2017.01823.

Stamatatos, L., Morris, L., Burton, D.R., and Mascola, J.R. (2009). Neutralizing antibodies generated during natural HIV-1 infection: good news for an HIV-1 vaccine? Nat Med 15, 866–870. 10.1038/nm.1949.

Stuart, T., Butler, A., Hoffman, P., Hafemeister, C., Papalexi, E., Mauck, W.M., Hao, Y., Stoeckius, M., Smibert, P., and Satija, R. (2019). Comprehensive Integration of Single-Cell Data. Cell 177, 1888–1902.e1821. 10.1016/j.cell.2019.05.031.

Subramanian, A., Tamayo, P., Mootha, V.K., Mukherjee, S., Ebert, B.L., Gillette, M.A., Paulovich, A., Pomeroy, S.L., Golub, T.R., Lander, E.S., and Mesirov, J.P. (2005). Gene set enrichment analysis: a knowledge-based approach for interpreting genome-wide expression profiles. Proc Natl Acad Sci U S A 102, 15545–15550. 10.1073/pnas.0506580102.

Sugawara, S., Thomas, D.L., and Balagopal, A. (2019). HIV-1 Infection and Type 1 Interferon: Navigating Through Uncertain Waters. AIDS Res Hum Retroviruses 35, 25–32. 10.1089/AID.2018.0161.

Takeuchi, O., and Akira, S. (2007). Recognition of viruses by innate immunity. Immunol Rev 220, 214–224. 10.1111/j.1600-065X.2007.00562.x.

Tan, L., Wang, Q., Zhang, D., Ding, J., Huang, Q., Tang, Y.Q., and Miao, H. (2020). Lymphopenia predicts disease severity of COVID-19: a descriptive and predictive study. Signal Transduct Target Ther 5, 33. 10.1038/s41392-020-0148-4.

Tang, F.C., Barbacioru, C., Wang, Y.Z., Nordman, E., Lee, C., Xu, N.L., Wang, X.H., Bodeau, J., Tuch, B.B., Siddiqui, A., et al. (2009). mRNA-Seq whole-transcriptome analysis of a single cell. Nature Methods 6, 377–U386. 10.1038/nmeth.1315.

Tangye, S.G., van de Weerdt, B.C.M., Avery, D.T., and Hodgkin, P.D. (2002). CD84 is up-regulated on a major population of human memory B cells and recruits the SH2 domain containing proteins SAP and EAT-2. European Journal of Immunology 32, 1640–1649. 10.1002/1521-4141(200206)32:6<1640::aid-immu1640>3.0.co;2-s.

Teijaro, J.R., Ng, C., Lee, A.M., Sullivan, B.M., Sheehan, K.C., Welch, M., Schreiber, R.D., de la Torre, J.C., and Oldstone, M.B. (2013). Persistent LCMV infection is controlled by blockade of type I interferon signaling. Science 340, 207–211. 10.1126/science.1235214.

Thangaraju, P., Venkatesan, N., Sudha, T.Y.S., Venkatesan, S., and Thangaraju, E. (2020). Role of Dupilumab in Approved Indications of COVID-19 Patient: an Efficacy-Based Nonsystematic Critical Analysis. SN Compr Clin Med, 1–5. 10.1007/s42399-020-00510-x.

Travaglini, K.J., Nabhan, A.N., Penland, L., Sinha, R., Gillich, A., Sit, R.V., Chang, S., Conley, S.D., Mori, Y., Seita, J., et al. (2020). A molecular cell atlas of the human lung from single-cell RNA sequencing. Nature 587. 10.1038/s41586-020-2922-4.

Treutlein, B., Brownfield, D.G., Wu, A.R., Neff, N.F., Mantalas, G.L., Espinoza, F.H., Desai, T.J., Krasnow, M.A., and Quake, S.R. (2014). Reconstructing lineage hierarchies of the distal lung epithelium using single-cell RNA-seq. Nature 509, 371-+. 10.1038/nature13173.

Trono, P., Tocci, A., Musella, M., Sistigu, A., and Nistico, P. (2021). Actin Cytoskeleton Dynamics and Type I IFN-Mediated Immune Response: A Dangerous Liaison in Cancer? Biology-Basel 10, 913. 10.3390/biology10090913.

Turner, J.S., O’Halloran, J.A., Kalaidina, E., Kim, W., Schmitz, A.J., Zhou, J.Q., Lei, T., Thapa, M., Chen, R.E., Case, J.B., et al. (2021). SARS-CoV-2 mRNA vaccines induce persistent human germinal centre responses. Nature 596, 109–113. 10.1038/s41586-021-03738-2.

Uematsu, S., and Akira, S. (2007). Toll-like receptors and Type I interferons. J Biol Chem 282, 15319–15323. 10.1074/jbc.R700009200.

Uhlen, M., Zhang, C., Lee, S., Sjostedt, E., Fagerberg, L., Bidkhori, G., Benfeitas, R., Arif, M., Liu, Z., Edfors, F., et al. (2017). A pathology atlas of the human cancer transcriptome. Science 357, 660-+. 10.1126/science.aan2507.

UNAIDS (2021). Fact Sheet - World AIDS Day 2021. UNAIDS.

Upasani, V., Rodenhuis-Zybert, I., and Cantaert, T. (2021). Antibody-independent functions of B cells during viral infections. Plos Pathogens 17, e1009708. 10.1371/journal.ppat.1009708.

Utay, N.S., and Douek, D.C. (2016). Interferons and HIV Infection: The Good, the Bad, and the Ugly. Pathog Immun 1, 107–116. 10.20411/pai.v1i1.125.

Wang, B., Kang, W., Zuo, J., and Sun, Y. (2017). The Significance of Type-I Interferons in the Pathogenesis and Therapy of Human Immunodeficiency Virus 1 Infection. Front Immunol 8, 1431. 10.3389/fimmu.2017.01431.

Wang, P., Meng, W., Han, S.-C., Li, C.-C., Wang, X.-J., and Wang, X.-J. (2016). The nucleolar protein GLTSCR2 is required for efficient viral replication. Scientific Reports 6, 36226. 10.1038/srep36226.

Wang, S., Zhang, Q., Hui, H., Agrawal, K., Karris, M.A.Y., and Rana, T.M. (2020). An atlas of immune cell exhaustion in HIV-infected individuals revealed by single-cell transcriptomics. Emerging Microbes & Infections 9, 2333–2347. 10.1080/22221751.2020.1826361.

Wang, Y., Chung, Y.R., Eitzinger, S., Palacio, N., Gregory, S., Bhattacharyya, M., and Penaloza-MacMaster, P. (2019). TLR4 signaling improves PD-1 blockade therapy during chronic viral infection. PLoS Pathog 15, e1007583. 10.1371/journal.ppat.1007583.

Ward-Kavanagh, L.K., Lin, W.W., Sedy, J.R., and Ware, C.F. (2016). The TNF Receptor Superfamily in Co-stimulating and Co-inhibitory Responses. Immunity 44, 1005–1019. 10.1016/j.immuni.2016.04.019.

Watanabe, R., Fujii, H., Shirai, T., Saito, S., Ishii, T., and Harigae, H. (2014). Autophagy plays a protective role as an anti-oxidant system in human T cells and represents a novel strategy for induction of T-cell apoptosis. European Journal of Immunology 44, 2508–2520. 10.1002/eji.201344248.

Weisberg, E., Parent, A., Yang, P.L., Sattler, M., Liu, Q., Liu, Q., Wang, J., Meng, C., Buhrlage, S.J., Gray, N., and Griffin, J.D. (2020). Repurposing of Kinase Inhibitors for Treatment of COVID-19. Pharmaceutical Research 37, 167. 10.1007/s11095-020-02851-7.

Wen, W., Su, W., Tang, H., Le, W., Zhang, X., Zheng, Y., Liu, X., Xie, L., Li, J., Ye, J., et al. (2020). Immune cell profiling of COVID-19 patients in the recovery stage by single-cell sequencing (vol 6, 31, 2020). Cell Discovery 6, 41. 10.1038/s41421-020-00187-5.

Wilk, A.J., Rustagi, A., Zhao, N.Q., Roque, J., Martinez-Colon, G.J., McKechnie, J.L., Ivison, G.T., Ranganath, T., Vergara, R., Hollis, T., et al. (2020). A single-cell atlas of the peripheral immune response in patients with severe COVID-19. Nature Medicine 26, 1070-+. 10.1038/s41591-020-0944-y.

Wilson, E.B., Yamada, D.H., Elsaesser, H., Herskovitz, J., Deng, J., Cheng, G., Aronow, B.J., Karp, C.L., and Brooks, D.G. (2013). Blockade of chronic type I interferon signaling to control persistent LCMV infection. Science 340, 202–207. 10.1126/science.1235208.

Wolf, F.A., Angerer, P., and Theis, F.J. (2018). SCANPY: large-scale single-cell gene expression data analysis. Genome Biol 19, 15. 10.1186/s13059-017-1382-0.

Wu, D., Harrison, D.L., Szasz, T., Yeh, C.F., Shentu, T.P., Meliton, A., Huang, R.T., Zhou, Z., Mutlu, G.M., Huang, J., and Fang, Y. (2021a). Single-cell metabolic imaging reveals a SLC2A3-dependent glycolytic burst in motile endothelial cells. Nat Metab 3, 714–727. 10.1038/s42255-021-00390-y.

Wu, J., Liang, B., Chen, C., Wang, H., Fang, Y., Shen, S., Yang, X., Wang, B., Chen, L., Chen, Q., et al. (2021b). SARS-CoV-2 infection induces sustained humoral immune responses in convalescent patients following symptomatic COVID-19. Nat Commun 12, 1813. 10.1038/s41467-021-22034-1.

Xu, C., Lopez, R., Mehlman, E., Regier, J., Jordan, M.I., and Yosef, N. (2021). Probabilistic harmonization and annotation of single-cell transcriptomics data with deep generative models. Molecular Systems Biology 17, e20209620. 10.15252/msb.20209620.

Xu, G., Qi, F., Li, H., Yang, Q., Wang, H., Wang, X., Liu, X., Zhao, J., Liao, X., Liu, Y., et al. (2020). The differential immune responses to COVID-19 in peripheral and lung revealed by single-cell RNA sequencing. Cell Discov 6, 73. 10.1038/s41421-020-00225-2.

Ye, L., Lee, J., Xu, L., Mohammed, A.U., Li, W., Hale, J.S., Tan, W.G., Wu, T., Davis, C.W., Ahmed, R., and Araki, K. (2017). mTOR Promotes Antiviral Humoral Immunity by Differentially Regulating CD4 Helper T Cell and B Cell Responses. J Virol 91. 10.1128/JVI.01653-16.

Yu, G., Wang, L.G., Han, Y., and He, Q.Y. (2012). clusterProfiler: an R package for comparing biological themes among gene clusters. OMICS 16, 284–287. 10.1089/omi.2011.0118.

Zhang, D., Guo, R., Lei, L., Liu, H., Wang, Y., Qian, H., Dai, T., Zhang, T., Lai, Y., Wang, J., et al. (2021). Frontline Science: COVID-19 infection induces readily detectable morphologic and inflammation-related phenotypic changes in peripheral blood monocytes. J Leukoc Biol 109, 13–22. 10.1002/JLB.4HI0720-470R.

Zhang, W., and Liu, H.T. (2002). MAPK signal pathways in the regulation of cell proliferation in mammalian cells. Cell Research 12, 9–18. 10.1038/sj.cr.7290105.

Zhao, L.-J., Hua, X., He, S.-F., Ren, H., and Qi, Z.-T. (2011). Interferon alpha regulates MAPK and STAT1 pathways in human hepatoma cells. Virology Journal 8, 157. 10.1186/1743-422x-8-157.

Zhu, L., Yang, P., Zhao, Y., Zhuang, Z., Wang, Z., Son, R., Zhang, J., Liu, C., Gao, Q., Xu, Q., et al. (2020). Single-Cell Sequencing of Peripheral Mononuclear Cells Reveals Distinct Immune Response Landscapes of COVID-19 and Influenza Patients. Immunity 53, 685-+. 10.1016/j.immuni.2020.07.009.

